# Characterization of immunosenescent alveolar macrophages in rhesus macaques

**DOI:** 10.1101/2025.10.10.681684

**Authors:** Jefferson G. C. Nagle, Raneesh Ramarapu, Laurent Zablocki-Thomas, Hai D. Nguyen, Dennis Hartigan-O’Connor, Amir Ardeshir, Elizabeth S. Didier, Woong-Ki Kim, Marcelo J. Kuroda

**Affiliations:** Department of Surgical and Radiological Sciences, School of Veterinary Medicine, University of California Davis, Davis, CA, 95616, USA; Department of Anatomy, Physiology & Cell Biology, School of Veterinary Medicine, University of California Davis, Davis, CA, 95616, USA; Division of Microbiology, Tulane National Primate Research Center, Covington, LA, 70433, USA; Department of Medical Microbiology and Immunology, School of Medicine, University of California Davis, Davis, CA, 95616, USA; Department of Microbiology & Immunology, Tulane University School of Medicine, New Orleans, LA, 70112, USA; Infectious Disease Unit, California National Primate Research Center, Davis, CA, 95616, USA

**Keywords:** aging, alveolar macrophages, inflammaging, rhesus macaques, single-cell RNA sequencing

## Abstract

As the global population ages, understanding the mechanisms underlying age-related diseases becomes increasingly important. Inflammaging, a state of chronic inflammation and immune dysregulation, is a key feature of aging. Macrophages, as master regulators of inflammation, are critical to this process. However, the effects of natural aging on alveolar macrophages (AMs) in the lungs remain poorly understood. In this study, we evaluated AM biology in older compared with young rhesus macaques. We compared cytokine profiles in plasma and bronchoalveolar lavage (BAL) fluid between young and aged macaques and performed single-cell RNA sequencing (scRNA-seq) to characterize age-related transcriptional and functional changes in AMs. Compared to young animals, AMs from aged animals exhibited similar cytokine inflammaging profile observed in the plasma of elderly humans. However, the general decrease of chemokine levels in BAL fluid relative to plasma suggest BAL may not reflect AM status in the lung tissues of rhesus macaques. scRNA-seq data corroborated plasma inflammaging findings, revealing increased inflammatory cytokine responses in AMs from older macaques. The consistent elevation of macrophage migration inhibitory factor (MIF) across plasma, AMs, and BAL fluid marks it as a promising biomarker and potential therapeutic target for age-associated inflammation. scRNA-seq further identified genes and pathways linked to macrophage senescence and turnover, suggesting potential targets for therapeutic intervention. Overall, our findings highlight the essential role of tissue-resident macrophages in pulmonary aging and underscores the need to further investigate their role in age-associated lung disease.

## Introduction

Rhesus macaques (*Macaca mulatta*) offer a well-established model for studying human aging due to their close physiological and genetic resemblance to humans (1,2). With a natural lifespan of over 25 years and a well-characterized immune system, rhesus macaques provide an ideal platform for investigating age-related changes in organ systems, including the lungs (3). In pulmonary research, this species provides the most valuable insight into both the natural progression of aging and the effects of pathogenic infections, such as simian immunodeficiency virus (SIV), which recapitulates many aspects of HIV infection in humans (3–6).

Aging in both humans and nonhuman primates (NHPs) is associated with progressive declines in lung function, structural remodeling of alveolar architecture, increased vulnerability to respiratory pathogens, and impaired resolution of inflammation (1,3). One of the central hallmarks of aging is “inflammaging”, a chronic, low-grade systemic inflammatory state that contributes to the onset and progression of multiple age-related diseases, including chronic obstructive pulmonary disease, idiopathic pulmonary fibrosis, and increased susceptibility to pneumonia (2,7). This persistent inflammatory milieu arises from altered immune cell function, increased cytokine production, and impaired resolution mechanisms.

Alveolar macrophages (AMs) comprise a long-lived and predominant immune cell population in the alveolar space that play essential roles in maintaining pulmonary homeostasis, clearing pathogens and debris, and orchestrating immune responses to environmental and infectious insults (8). However, the extent to which natural aging alters AM phenotype, function, and gene expression remains poorly understood. Prior studies in mice indicate that AMs undergo age-related changes in polarization, phagocytosis, and response to stimuli (9). In humans, age-associated transcriptomic changes have been identified in peripheral monocytes and tissue macrophages (10), yet the aging trajectory of AMs specifically has not been fully characterized in human and translationally relevant models like rhesus macaques.

Emerging evidence suggests that AMs not only respond to systemic inflammatory cues but may also contribute directly to the inflammaging phenotype (8,11). Their longevity, self-renewal capacity, and residence in a uniquely exposed environment position them as critical regulators of both local and systemic immune responses during aging. However, the molecular and functional characteristics of AMs in naturally aged NHPs remain largely unexplored. To address this gap, we investigated the biology of AMs in young and aged rhesus macaques under steady-state conditions. We examined multiplex cytokine assays to compare inflammatory profiles in plasma and bronchoalveolar lavage (BAL) fluid, and single-cell RNA sequencing (scRNA-seq) to define transcriptional changes associated with AM aging. By integrating functional, molecular, and immunohistochemical analyses, we aimed to (*i*) determine whether AMs undergo age-related phenotypic and transcriptional reprogramming, (*ii*) assess how aging affects cytokine dynamics across compartments, and (*iii*) identify candidate genes and pathways involved in pulmonary inflammaging. This integrative approach provides novel insights into the role of AMs in lung aging and supports the hypothesis that the long-lived AMs become transcriptionally reprogrammed to support a pro-inflammatory and metabolically altered state. These findings lay the groundwork for future studies targeting macrophage-mediated immune dysregulation in age-associated lung diseases.

## Material and Methods

### Animal work approval

All animal work was performed at the University of California, Davis, an institution accredited by the Association for Assessment and Accreditation of Laboratory Animal Care International (AAALAC). Animal care adhered to the 2011 *Guide for the Care and Use of Laboratory Animals* provided by the Institute for Laboratory Animal Research (12). The study was approved by the Institutional Animal Care and Use Committee (IACUC) at the University of California, Davis, under protocols 22107, 22399, and 22484.

### Animal model

A total of 144 Indian rhesus macaques (*Macaca mulatta*) were used for this study. The young group consisted of 62 rhesus macaques aged 5–12 years, while the old group included 82 rhesus macaques aged 15–24 years. All animals were socially housed at the California National Primate Research Center (CNPRC) and were specific pathogen-free for SIV, simian type D retrovirus, and simian T-cell leukemia virus type 1 at the time of enrollment.

Animals were provided with water *ad libitum* and received commercial high-protein diet (Ralston Purina Co.) supplemented with fresh fruits. Euthanasia was performed humanely by veterinarians, either at pre-defined experimental endpoints or based on clinical indicators (e.g., weight loss, poor body condition, lethargy, or appetite loss), in compliance with IACUC guidelines and national standards. The procedure included anesthesia with ketamine hydrocholoride (HCl) (10 mg/kg) followed by an overdose of sodium pentobarbital, consistent with the recommendations of the *American Veterinary Medical Association Guidelines on Euthanasia* (13). Demographic data for the animals are presented in **Error! Reference source not found.Error! Reference source not found.**, and an overview of the study design is shown in **Figure 1**. All possible measures were taken to minimize animal discomfort. For routine procedures, such as blood collection and physical examinations, animals were fully anesthetized with ketamine HCl under the direction of a veterinarian.

**Figure 1.**
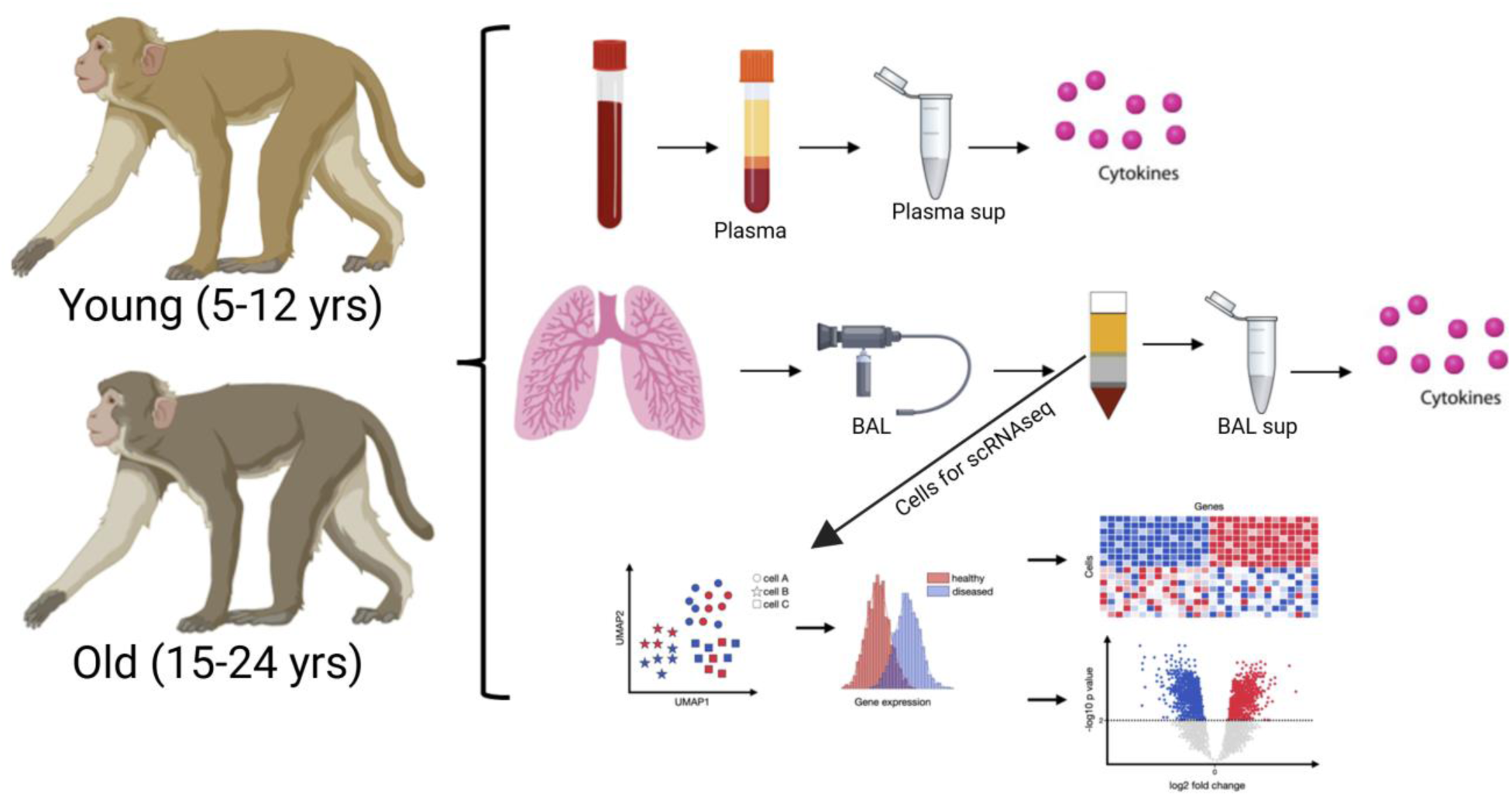
Experimental Design. Plasma aliquots from both groups of animals were saved at -80 °C for cytokine testing. Lungs were collected from euthanized animals. The left lower lobes were dissected for bronchoalveolar lavage; supernatants were isolated and saved at -80 °C for cytokine testing. The upper left lobes were instilled with 10% formalin solution for paraffin block sections. The right-side lobes were collected in R10 media solution and digested for isolation of alveolar macrophages. One million cells were separated for flowcytometry immunostaining and 1x10^5^ cells were separated for scRNA-seq.

### Lung collection

At necropsy, the lung tissue of each animal was removed from the thoracic cavity and the different lobes further separated for different purposes. Rhesus macaque lungs are anatomically divided into right and left lungs, which are further divided into lobes as shown in **Error! Reference source not found.**.

### Bronchoalveolar lavage collection and alveolar macrophage isolation

BAL samples were centrifuged at 673 g for 5 min (Eppendorf 5810R Refrigerated Centrifuge, Hamburg, Germany). BAL supernatants were aliquoted into 1.5-mL tubes and stored at -80°C for cytokine assays. Cell pellets were resuspended with 5 mL of 2% PBS-FBS (PBS containing 2% FBS) and filtered through a 100-µm cell strainer (Falcon™, Catalog No. 08-771-19). Tubes were rinsed with an extra 5 mL of 2% PBS-FBS and filtered with 100-µm cell strainer. Cells were centrifuged at 673 g for 5 min and resuspended in red blood cell (RBC) Ammonium-Chloride-Potassium (ACK) Lysing Buffer (Gibco™, Catalog No. A1049201) for 10 min at room temperature. Ten mL of 2% PBS-FBS was added followed by centrifugation at 673 g for 5 min. The cells then were resuspended in 5 ml of 37°C pre-warmed in RPMI 1640 Medium, GlutaMAX (Gibco™; cat. no. 61870036, Grand Island, NY, USA) supplemented with 10% FBS (Gibco™; cat no. 10437-028; Grand Island, NY, USA), 100 IU/ml of penicillin/streptomycin (MP Biomedicals, Inc; cat no. ICN1670249; Santa Ana, CA, USA), 25mM of HEPES (MP Biomedicals, Inc; cat no. ICN1688449; Santa Ana, CA, USA). After counting by trypan blue exclusion, aliquots of live cells were adjusted to 1 x 10^6^ cells for immunostaining and flow cytometry collection and the rest of the cells were centrifuged at 673 g for 5 min and suspended in BAMBANKER™ serum-free cell culture freezing medium (Wako Laboratory Chemicals, cat no. 302-14681; Richmond, VA, USA) and stored in liquid nitrogen for scRNA-seq and further assays.

### Multiplex cytokine and chemokine analysis

BAL supernatants and plasma were tested using Cytokine 29-Plex Monkey Panel and Luminex platform technology (Novex; Life Technologies, cat# LPC0005M) according to the instructions of the manufacturer. **Error! Reference source not found.** has the results of exact p-value and adjusted p-value with Bonferroni correction to counteract the multiple comparisons problem and Benjamini-Hochberg Procedure for false discovery rate correction (FDR) for plasma and **Error! Reference source not found.** has the results for BAL supernatant. Average group data are given as mean ± SEM or as median and range (minimum – maximum). Significance of differences between groups was calculated using the nonparametric *t* test (Mann–Whitney *U* test). Briefly, 50 μL of each sample was incubated in duplicate for 2 h with anti-cytokine/chemokine Ab–coated beads. All samples were incubated on the same plate at room temperature on an orbital shaker at 600 rpm. The cytokine/chemokine standard curve samples provided with the kit were included on the plate. After washing, biotinylated Abs supplied with the kit were added to the beads and incubated for 1 h at room temperature on an orbital shaker at 600 rpm and after another washing step, streptavidin–R-PE was added and incubated for 30 min at room temperature on an orbital shaker at 600 rpm. After removal of the streptavidin–R-PE and washing, the beads were resuspended in wash solution, after which the plate was read on a Bio-Plex 200 system (Bio-Rad, Veenendaal, the Netherlands). Concentrations of the cytokines and chemokines in the samples were calculated using the Bio-Plex Manager Software v.4.1 program (Bio-Rad Laboratories, Hercules, CA, USA). Data analyses were performed with R programing language (version 4.4.1) (14) and RStudio software (version “Cranberry Hibiscus”, Build 394) (15). Multiple packages were utilized during analysis including tidyverse (16), ggplot2 (17,18), and dplyr (19) packages were utilized for data manipulation and visualization. The statistical analyses and preparation of cytokine comparison plots were performed using boxplots, with significance annotations based on statistical tests. The FC values were derived from the log-transformed cytokine values across different groups (Young vs. Old) in both plasma and BAL supernatant samples.

Due to the skewed distribution of cytokine values, as determined by the Shapiro-Wilk normality test, non-parametric tests were applied. The Mann-Whitney U test was used to evaluate differences between groups, and *p*-values were adjusted using the Bonferroni correction to counteract the multiple comparisons problem and Benjamini-Hochberg Procedure for False Discovery Rate (FDR) method to account for multiple comparisons. *P*-values were denoted with the following significance symbols: p < 0.001 (\*\*\**), p < 0.01 (**), and p < 0.05 (*\*), with ‘NS’ used to represent non-significant differences. Cytokine levels that were below detection limits were excluded from the analysis. Outliers were identified and removed using the interquartile range (IQR) method, where data points falling outside 1.5 times the IQR were considered outliers. An adjusted p-value of less than 0.05 was considered statistically significant for all comparisons.

### Single Cell RNA sequence workflow

#### Cell preparation

Cryopreserved BAL isolated alveolar macrophages were rapidly thawed in a 37°C water bath until pea-sized frozen cryopreserved cells remained. Cells were resuspended by gradually adding 13 mL of 37°C pre-warmed RPMI 1640 Medium with GlutaMAX™ (Gibco™, Cat. #61870036, Grand Island, NY, USA), supplemented with 10% FBS (Gibco™, Cat. #10437-028), 100 IU/mL penicillin/streptomycin (MP Biomedicals, Cat. #ICN1670249, Santa Ana, CA, USA), and 25 mM HEPES (MP Biomedicals, Cat. #ICN1688449, Santa Ana, CA, USA). Cells were centrifuged at 300 x g for 10 min at 4°C (Eppendorf 5810R Refrigerated Centrifuge, Hamburg, Germany). Before centrifugation, 20 µL of the cell suspension were removed for manual cell counting using a hemocytometer.

#### Dead cell removal

AMs were processed using the MACS Miltenyi Biotec Dead Cell Removal Kit (Cat. #130-090-101) following the manufacturer’s instructions. Cells were centrifuged at 300 x g for 5 min, and the supernatant was removed. The cell pellet was resuspended in 100 µL of Dead Cell Removal MicroBeads (per 10^7^ total cells), incubated at room temperature for 15 min, and passed through MS columns (Cat. #130-042-201) on an OctoMACS separator (Cat. #130-042-109). The effluent containing live cells was collected, centrifuged again at 300 x g for 5 min, and resuspended in cold 0.5% Bovine Serum Albumin (BSA, Sigma-Aldrich, Cat. #A8412) in PBS (Corning™, Cat. #MT21040CV, Corning, NY, USA). Final live cell counts were manually performed using a hemocytometer.

#### Fixation and permeabilization

The cells were fixed and permeabilized using the Parse Biosciences Fixation Kit (Seattle, WA, USA) according to the manufacturer’s protocol. After centrifugation at 300 x g for 10 min at 4°C, cells were resuspended in cold cell buffer and filtered through a 40-µm strainer. Fixation Solution (250 µL) was immediately added, mixed by pipetting, and incubated on ice for 10 min. Then, 80 µL of Permeabilization Solution were added followed by incubation for 3 min on ice and then addition of 4 mL of cold Neutralization Buffer. After centrifugation, cells were resuspended in 300 µL of cold cell buffer and filtered again through a 40-µm strainer. Final cell counts were manually performed prior to cryopreservation.

#### Freezing fixed sample

For samples containing more than 500,000 cells, 5 µL of DMSO were added, mixed gently, and incubated for 1 min on ice. This process was repeated twice to ensure a final DMSO volume of 15 µL. The cell suspension was pipetted thoroughly to ensure even distribution of DMSO before aliquoting into 2 x 10^5^ cells per sample for cryopreservation.

#### Data Acquisition and Processing

A total of 100,000 cells from 24 donors was used, resulting in an average of approximately 4,166 cells per donor. These cells were subjected to the Single Cell Whole Transcriptome Kit v2 (Parse Biosciences, Seattle, WA, USA) for library construction. The scRNA-seq libraries for Parse Biosciences platform were sequenced using the Illumina NovaSeq 6000 platform.

After sequencing, raw data were processed using the Parse Biosciences pipeline, which included demultiplexing, alignment, and UMI counting. The sequencing reads were mapped to the MMul_10 reference genome (assembly GCF_003339765.1) using NCB1 Gene Build 103 for annotation. The resulting cell x barcode matrices were further analyzed using the *Seurat* package (v5.1.0) (20) for downstream scRNA-seq analysis. Quality control filters were applied to remove samples with less than 100 numbers of features (genes) per cell, and less than 3000 total RNA content per cell.

Gene expression matrices were loaded into the R package *Seurat* for quality control and downstream analyses (21). Quality control parameters were applied based on the best practices for scRNA-seq (22–24). We removed cells fulfilling any of the following criteria: (1) lowest number of unique genes detected in any cell (nFeature_RNA) > 200. A very low number of detected genes suggest poor-quality or dying cells with degraded RNA. (2) maximum number of unique genes detected in any cell (nFeature_RNA) < 4,500. A high number of detected genes suggest doublets. (3) number of minimal UMI count for RNA molecules in a cell (nCount_RNA) > 500. Very low UMI counts often indicate poor quality cells or empty droplets; (4) number of maximal UMI count for RNA molecules in a cell (nCount_RNA) < 20,000. It is important to remove cells with extremely high UMI counts, as they may represent doublets or overly high sequencing depth (outliers); and (5) percentage of mitochondrial genes in any cell (percent.mt) < 5. By removing cells with more than 5% of mitochondrial genes, we ensure removal of dying cells.

Cell doublets were detected and removed using the R package *DoubletFinder* (2.0.3) (25). Genes expressed in less than 8 cells were discarded. Gene counts were then normalized and transformed with *Seurat::NormalizeData()*, which normalizes the total counts for each cell by the total counts, multiplies this by a scale factor of 10,000, and log-transforms the result.

#### Single cell data analysis quality control

The scRNA-seq data were processed, resulting in a total of 8,511 cells. Key quality control metrics included a high base call quality, with bc1_Q30, bc2_Q30, and bc3_Q30 scores of 0.889, 0.911, and 0.923, respectively. The cDNA Q30 score was 0.9044, indicating excellent read quality across the libraries. On average, 1,315 genes were detected per cell, with a median of 1,549.6 transcripts per cell. The mean number of reads per cell was 74,331.54, supporting robust transcriptomic coverage for individual cells. In total, 44,562,775 transcripts and 25,312 genes were detected across all cells, indicating a comprehensive capture of the transcriptome.

The alignment statistics revealed that 435,382,521 reads were successfully aligned, with 321,257,629 of these being uniquely mapped. A relatively small proportion, 20,391,757 reads, were found to be multimapping, ensuring high confidence in the alignment process. The fraction of reads mapped to the transcriptome was 61.33%, indicating efficient capture of coding regions. Sequencing saturation reached 83.31%, and the valid barcode fraction was 68.82%, both metrics indicating effective utilization of sequencing resources and sample quality. PolyN Q30, a measure of the poly-N tail quality, was 0.919, further underscoring the high quality of the sequencing data.

#### Single-Cell Data Analysis Workflow

*Seurat* v5 was utilized for downstream analysis and was applied according to the package vignettes (26). Filtered feature matrices were independently normalized and scaled using the function *SCTransform* v2 (glmGamPoi) (27). Using the elbow plot method, we determined that the first 10 principal components captured the majority of variance in the data and were optimal for guiding further analysis, including clustering and visualization (**Error! Reference source not found.**).

#### Clustering Analysis

The *clustree* analysis revealed that a resolution of 0.1 provided the best balance between clear cluster delineation and biological interpretability (**Error! Reference source not found.**) (28). At this resolution, clusters appeared well-separated with minimal overlap, indicating that they likely represent transcriptionally distinct cell populations.

To further optimize clustering performance, we systematically tested different combinations of dimensional reduction parameters and gene thresholds. The best-performing combination was 10 PCs and resolution = 0.1, yielding the highest silhouette score of 0.1631 (**Error! Reference source not found.**). We also compared nine thresholds for the number of genes used in clustering, ranging from 2,000 to 15,000 genes derived from the SCT assay, and found that selecting 3,000 highly variable genes resulted in the best silhouette score of 0.1642 (**Error! Reference source not found.**). These analyses confirmed that resolution = 0.1 and 3,000 variable genes captured the optimal structure of alveolar macrophage heterogeneity in our dataset. (**Error! Reference source not found.**).

#### Identifying cell clusters

To focus our analysis on natural aging, we stratified the scRNA-seq dataset excluding any animals or samples associated with experimental interventions or disease models. The final dataset included 42,147 alveolar cells derived from 7 young (5–12 years) and 5 old (15–24 years) rhesus macaques. Of these, 25,396 cells originated from young animals and 16,751 from old animals (**Figure 5A**).

### Statistical Analyses

All statistical analyses and graph preparations were performed using R programing language (version 4.4.1, R Core Team, 2024) (14) and RStudio software (version “Cranberry Hibiscus”, Build 394) (15). Additional packages used were dependent on the specific analysis. *Seurat* package (v5.1.0) (20) was used for downstream scRNA-seq analysis. Differential expression analysis was conducted using the FindMarkers function. Shapiro-Wilk normality test was applied, and non-normal distribution was confirmed for all samples. Consequently, Mann– Whitney U test was applied to evaluate differences between two study groups. Differential expression analysis (DEA) of AMs from young and aged groups was conducted using *MAST* (v1.16.0). Genes with an adjusted p-value < 0.05 and an absolute FC ≥ 2 were considered significant and subsequently analyzed for functional enrichment using PANTHER Gene Ontology (GO) and Ingenuity Pathway Analysis (IPA). Additional insights into pathway enrichment were obtained through KEGG and targeted GO analyses.

## Results

### Systemic Cytokine Levels Increase with Age in Rhesus Macaques

Aging in rhesus macaques mirrors human “inflammaging” and is associated with elevated systemic levels of pro-inflammatory cytokines (29). Older animals (15–24 years) displayed significantly increased plasma concentrations of IL-1β, IL-2, IL-6, IL-12, IL-15, IFN-γ, TNF-α, and G-CSF compared to younger macaques (5–12 years), with IL-1β, IL-2, and TNF-α showing the most pronounced elevations (*p* < 0.001; **Figure 2**). The anti-inflammatory cytokine IL-1RA was also significantly elevated in older animals (*p* < 0.001), while IL-10 levels remained unchanged (*p* = 0.811), suggesting selective regulation of inflammatory mediators during aging.

**Figure 2.**
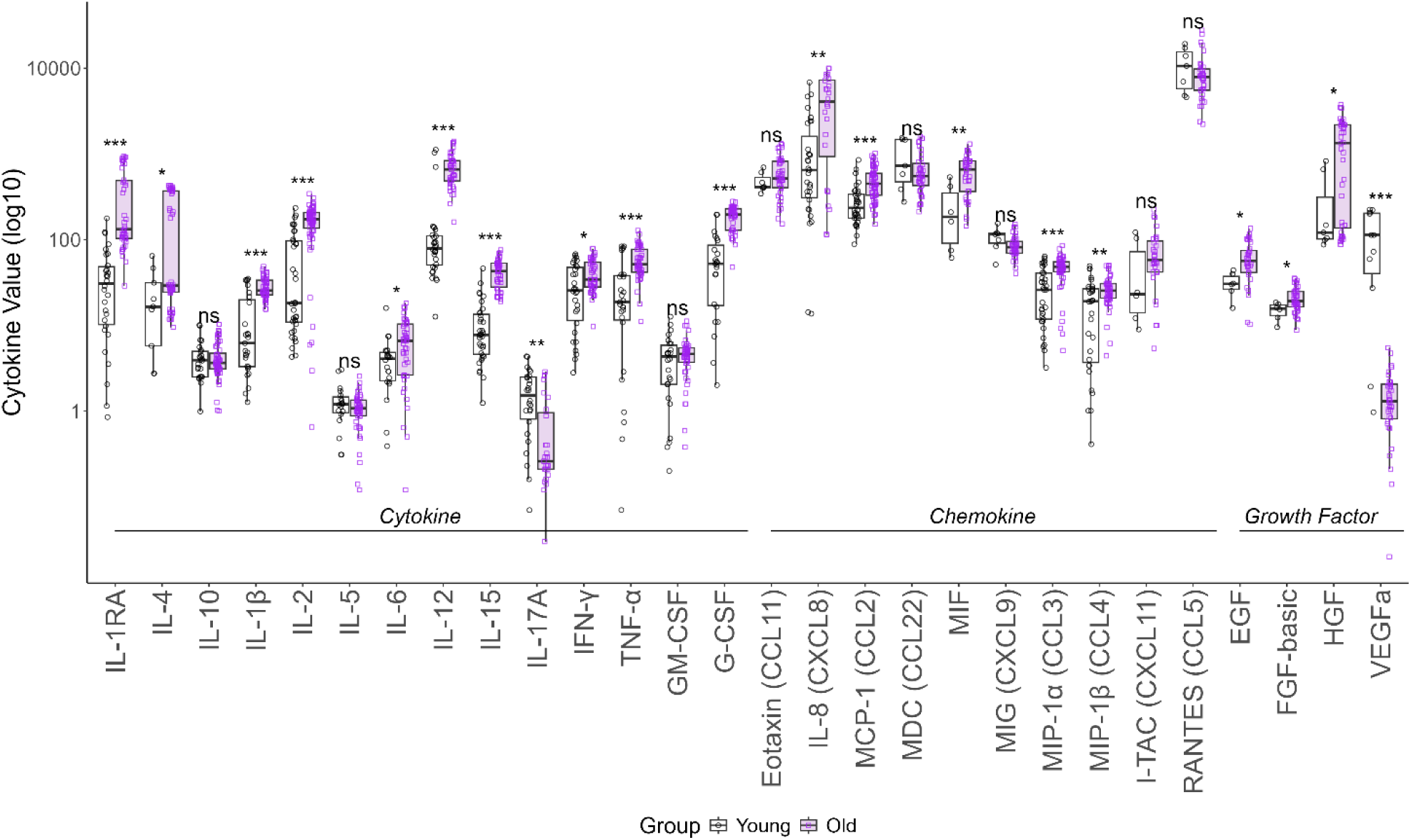
Elevated Plasma Cytokines Reflect Systemic Inflammaging During Natural Aging. Plasma cytokine levels, originally measured in pg/mL via a multiplex immunoassay, were log10-transformed to improve distributional properties and comparability across analytes. Cytokine levels were compared between young (5-12 years) and old (15-24 years) groups using the Mann-Whitney U test, as most cytokines (27 out of 30) did not meet normality assumptions (Shapiro-Wilk test, p < 0.05). MIF, EGF, and FGF-basic, despite being normally distributed, were analyzed using the non-parametric test for consistency. False Discovery Rate (FDR) correction was applied to account for multiple comparisons. Significant differences were found in IL-17A and VEGFa, both elevated in young animals. Several cytokines, including IL-10, IL-5, GM-CSF, Eotaxin (CCL11), MDC (CCL22), MIG (CXCL9), I-TAC (CXCL11), and RANTES (CCL5), showed no significant differences across age groups. All other cytokines showed a significant increase with aging.

In addition to cytokines, older macaques exhibited significantly higher levels of multiple chemokines. These included IL-8 (CXCL8), MIF, and MIP-1β (*p* < 0.01), as well as MCP-1 (CCL2), MIP-1α (CCL3) (*p* < 0.001), and I-TAC (CXCL11) (*p* < 0.05), as confirmed by Mann-Whitney U testing. These chemokine elevations may reflect enhanced leukocyte recruitment signaling with age.

Growth factors including EGF, FGF-basic, and HGF were also significantly elevated in older animals (*p* < 0.05), suggesting age-associated alterations in tissue repair and angiogenesis pathways. In contrast, IL-17A (*p* < 0.01) and VEGFa (*p* < 0.001) were elevated in younger animals, indicating selective immune and vascular changes that decline with age. These opposing trends highlight the complexity of immune modulation over the lifespan.

Collectively, these findings confirm that rhesus macaques display systemic age-associated increases in pro-inflammatory cytokines, chemokines, and growth factors, consistent with human inflammaging. The observed systemic immune shifts prompted further investigation into AMs as key contributors to the aging inflammatory phenotype.

### Alveolar Macrophages Mirror Systemic Inflammatory Changes

To evaluate the contribution of AMs to systemic inflammation, cytokine gene expression was analyzed in AMs isolated from BAL fluid using scRNA-seq. This approach allowed us to assess cytokine production at single-cell resolution in a tissue-resident immune population. By focusing on AMs, we aimed to determine whether their transcriptional profile reflected the systemic inflammatory trends observed in plasma.

Aged AMs demonstrated increased expression of key pro-inflammatory mediators, including IL-1β, IL-5, IL-6, TNF-α, GM-CSF, IL-8 (CXCL8), MIF, MIP-1α (CCL3), RANTES (CCL5), IP-10, EGF, and VEGFa (**Figure 3**). These patterns mirrored the elevated levels of corresponding cytokines in plasma, indicating that AMs may be important contributors to systemic inflammaging. The consistent upregulation across both compartments supports the hypothesis that aging impacts immune signaling in both circulating and tissue-resident immune cells. Conversely, several cytokines and chemokines were downregulated in aged AMs, including IL-1RA, IL-4, IL-15, IFN-γ, G-CSF, Eotaxin, MCP-1 (CCL2), MDC (CCL22), MIG (CXCL9), I-TAC (CXCL11), and HGF (**Figure 3**).

**Figure 3.**
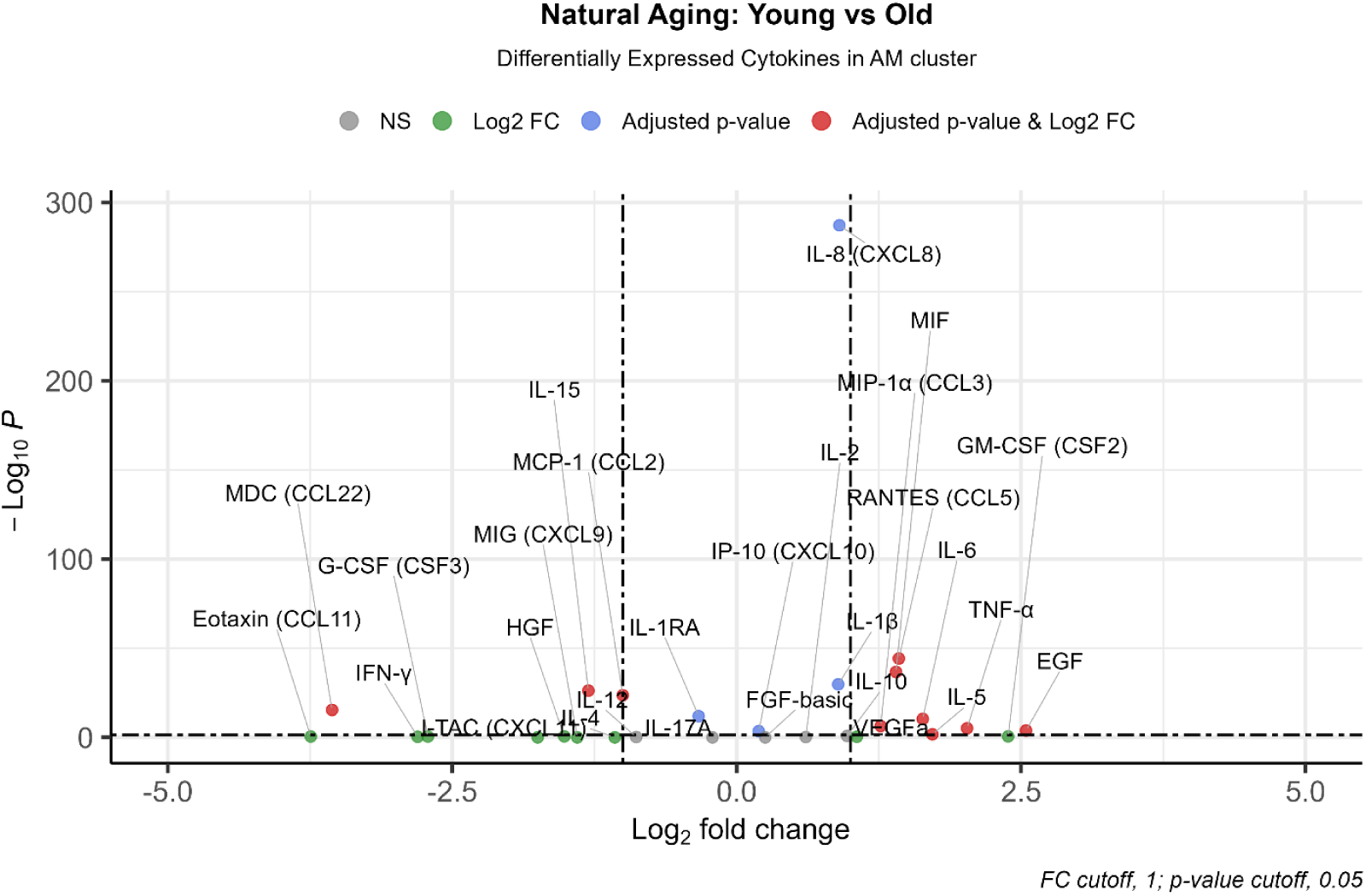
Alveolar Macrophage Cytokine Expression Highlights Pro-Inflammatory Shifts During Natural Aging. Volcano plot depicting cytokine gene expression changes in alveolar macrophages (AMs) during natural aging between young (n= 7) and old (n= 5). Values below 0 of Log2 fold are downregulated and above 0 are upregulated in the old group. Extremely significant upregulation (Log2-Fold >1 & p-value adjust < 0.05) includes pro-inflammatory cytokines, IL-5, IL-6, TNF-α, chemokine MIF, MIP-1α (CCL3), RANTES (CCL5) and growth factor EGF. Significant upregulation (Log2-Fold >1 OR p-value adjust < 0.05) includes IL-1β, GM-CSF, IL-8 (CXCL8), IP-10, and growth factor VEGFa. Extremely significant downregulation in the old (Log2-Fold > -1 & p-value adjust < 0.05) are IL-15, MCP-1 (CCL2), and MDC (CCL22). Significant downregulation in the old (Log2-Fold > -1 OR p-value adjust < 0.05) includes IL-1RA, IL-4, IFN-γ, G-CSF, Eotaxin (CCL11), MIG (CXCL9) I-TAC (CXCL11) and HGF. As expected, we observed in old macaques, a shift towards a heightened inflammatory state coupled with a decline in anti-inflammatory response. Notably, MIF emerges as consistently elevated, reinforcing its central role in inflammaging. NS (non-significant); Log2 FC (twice increased or decreased in the comparison group in comparison with the control group); Adjusted p-value (p-value corrected with Bonferroni to counteract the multiple comparisons); Adjusted p-value & Log2FC (Extremely significant cytokines since they are twice increased and decreased in old vs young and has a p-value adjust < 0.05). *The MIP-1b (CCL4) gene was not found in AM scRNA-seq*.

### Cytokine Levels in BAL Supernatant Do Not Fully Reflect Plasma or AM Trends

Given the pronounced age-related increases in cytokines within plasma and their alignment with AM gene expression (9), we next analyzed cytokine concentrations in BAL supernatant. BAL fluid is often used as a minimally invasive method to assess pulmonary immune responses, making it an attractive candidate for aging-related biomarker studies. However, our results revealed a contrasting pattern in cytokine levels compared to those observed in plasma and AMs. In aged animals, the majority of cytokines measured in BAL supernatant were decreased relative to younger animals (**Figure 4**). Only IL-17A, MIF, and VEGFa exhibited consistent directional changes across all compartments (plasma, BAL supernatant, and AM gene expression). Notably, MIF was the only cytokine elevated in all three sources, reinforcing its potential central role in the inflammatory processes associated with aging. The divergence between plasma and BAL supernatant cytokine levels suggests that the latter may be influenced by cell types other than AMs, based on the detection of epithelial cells, neutrophils, and other immune populations residing within the alveolar space (**Error! Reference source not found.**).

**Figure 4.**
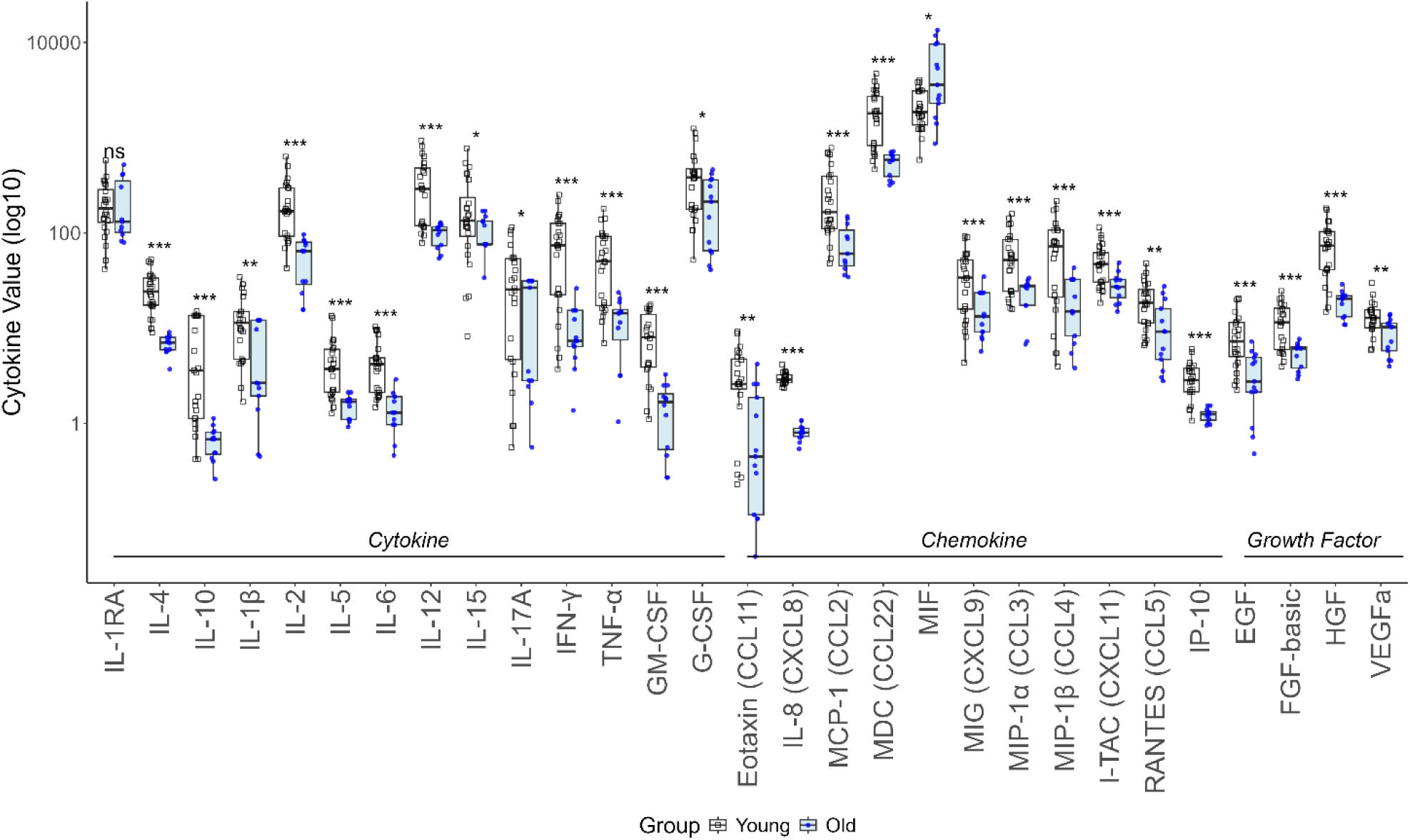
BAL Supernatant Cytokine Levels Decrease with Age Despite Systemic Inflammaging. BAL supernatant cytokine levels were compared between young (5-12 years) and old (15-24 years) groups. The Mann-Whitney U test was applied, as most cytokines did not meet normality assumptions (Shapiro-Wilk test, p < 0.05). p-value adjusted correction was done using Bonferroni to counteract the multiple comparisons problem and Benjamini-Hochberg Procedure for false discovery rate correction (FDR). IL-1RA was the only non-significant cytokine. Significant age-related differences were observed in several cytokines, with MIF being the only one that increased with aging as seen in plasma. All other cytokines showed age-related differences with significant increase in young animals compared to aged animals.

### Single-Cell RNA Sequencing Analysis of Alveolar Macrophages in Young and Old Rhesus Macaques

Given the observed elevation of pro-inflammatory cytokines in plasma during natural aging and the proposed contribution of AMs to this systemic inflammatory profile, we next sought to investigate the cellular and molecular underpinnings of these age-associated changes. To accomplish this, we performed scRNA-seq on AMs isolated from BAL fluid of 7 young (5–12 years) and 5 old (15–24 years) rhesus macaques. This approach allowed us to examine cellular transcriptional heterogeneity in detail and address two key questions: (*i*) How do age-related changes in AM gene expression correspond to the systemic cytokine trends observed in plasma? and (*ii*) What functional pathways change in AMs during natural aging?

By leveraging scRNA-seq, we were able to characterize the transcriptional landscape of alveolar macrophages at single-cell resolution. This analysis enables the identification of gene expression changes associated with aging and provided insights into the cellular pathways that contribute to inflammaging. The scRNA-seq platform offered an unbiased means to interrogate molecular aging in tissue-resident immune cells and to uncover functional programs that may underlie age-related immune dysregulation. We applied the previously optimized clustering parameters, 3,000 highly variable genes, 10 principal components, and a resolution of 0.1, to ensure robust and biologically meaningful cluster identification. Dimensionality reduction via Uniform Manifold Approximation and Projection (UMAP) revealed five transcriptionally distinct cell clusters in the lung alveolar compartment (**Figure 5B**). These clusters represented the major immune and stromal cell types present in BAL fluid under steady-state aging conditions. All clusters present in the data were well integrated between both young and old animals (**Figure 5C**). AMs compromise the vast majority of the cells isolated from BAL in both groups (**Figure 5D**).

**Figure 5.**
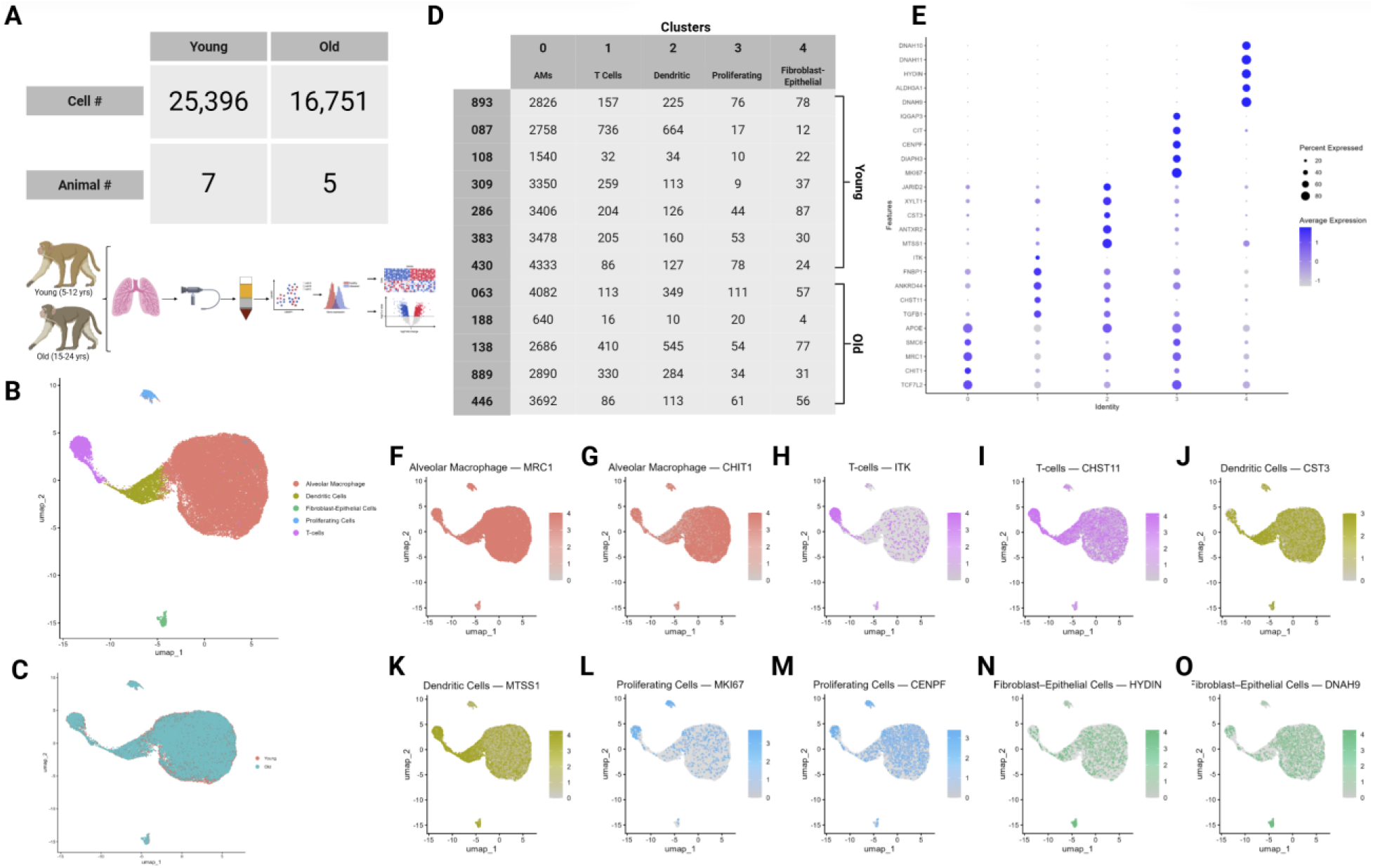
Alveolar Macrophages Defined the Largest Clusters in BAL Cells. scRNA-seq analysis of BAL cell isolates revealed 5 major cell types across 7 young and 5 old rhesus macaques. (A) Table of cell counts from each age group. (B) UMAP plot demonstrating the clustering results for immune cells from all 12 healthy macaques of different age groups. (C) UMAP plot of global clustering colored by age. (D) Table of cell type counts across clusters by age. (E) Dot plot demonstrating cell type specific marker expression of the 5 major cell types. (F–O) Feature UMAP plots demonstrating expression profile of key markers for each of the 5 different cell types.

Cluster annotation was based on the top five marker genes in each group (**Figure 5E**) cross-referenced with published cell-type–specific gene signatures (30). The identified clusters were classified as follows: cluster 0 (AMs), cluster 1 (T cells), cluster 2 (dendritic cells), cluster 3 (proliferating cells), and cluster 4 (fibroblast/epithelial cells) (**Figure 5B)**. Feature UMAP plots demonstrate expression profile of key markers for each of the 5 distinct cell types identified (**Figure 5F-O)**. This clustering framework provided the basis for subsequent analyses of age-associated transcriptional changes in AMs.

### Alveolar Macrophages Exhibit Distinct Immunometabolic and Inflammatory Signatures with Age

Our scRNA-seq analysis identified a transcriptionally distinct AM cluster characterized by a unique and multifunctional gene expression profile. This cluster exhibited the highest expression of canonical macrophage markers and genes associated with tissue maintenance, innate immune responses, and lipid metabolism (**Figure 5E**). The transcriptional complexity within this population highlights the functional heterogeneity of AMs in the aging lung microenvironment.

Notably, the AM cluster expressed elevated levels of *TCF7L2* and *MYO1F*, genes implicated in transcriptional regulation and macrophage-specific activity, respectively (31,32). Additional markers such as *TMSB4X*, *ACAD10*, *B2M*, *SNORD89*, and *APOE* were also highly expressed, reflecting roles in cytoskeletal dynamics, metabolic regulation, and antigen presentation (33–36). This profile underscores the importance of AMs in orchestrating immune surveillance and metabolic adaptation within the lung.

The macrophage mannose receptor *MRC1* (also known as CD206), a hallmark of tissue-resident alveolar macrophages, was prominently expressed in this cluster, confirming identity of AMs. CD206 plays critical roles in endocytosis, tissue remodeling, and immune modulation (37,38). Other highly enriched genes included *DDHD1*, *TRPM2*, *IL1B*, *CCL18*, *CXCL8*, *CD68*, and *CHIT1*, all of which are associated with lipid signaling, chemokine production, and inflammatory responses. These findings provide strong evidence that AMs in aging lungs retain multifunctional roles essential for pulmonary homeostasis, host defense, and immune regulation (36,39,40).

### Distinct Transcriptional Profiles of T-Cell and Dendritic Cell Populations in the Lung

Cluster 1 identified T-cells based on elevated expression of several key markers associated with T-cell function, development, activation, and exhaustion (**Figure 5E**). These markers include SKAP1, ETS1, ITK, TOX, CHST11, BICDL1, and CAMK4, all well-known for their critical roles in T-cell biology. SKAP1 (Src Kinase Associated Phosphoprotein 1) is an adaptor protein that plays a crucial role in T-cell receptor (TCR) signaling by enhancing the MAP kinase pathway, thereby promoting T-cell activation (41,42). Additionally, SKAP1 facilitates optimal conjugation between T-cells and antigen-presenting cells by clustering integrin ITGAL (LFA-1) on the T-cell surface, which is essential for effective immune synapse formation (43,44). ETS1 (ETS Proto-Oncogene 1) is a transcription factor vital for T-cell development and function. It regulates genes necessary for T-cell activation and differentiation, with studies demonstrating its essential role in maintaining proper immune responses (45,46). ITK (IL2-Inducible T-Cell Kinase) is a tyrosine kinase predominantly expressed in T cells that mediates TCR signaling by activating PLCγ, which generates second messengers to propagate activation signals (47). Proper ITK function is critical for T-cell development and effective immune responses (48). TOX (Thymocyte Selection-Associated High Mobility Group Box) is a transcription factor implicated as a key regulator of T-cell exhaustion, a dysfunctional state seen in chronic infections and cancer. Elevated TOX expression drives epigenetic remodeling, leading to increased inhibitory receptor expression and reduced effector functions in exhausted T cells (49).

Cluster 2 comprises dendritic cells, characterized by elevated expression of several key markers including CD86, CPVL, GPR183, MTSS1, and ANTXR2 (**Figure 5E**). CD86 (Cluster of Differentiation 86) is a co-stimulatory molecule constitutively expressed on dendritic cells and other antigen-presenting cells, providing essential signals for T-cell activation and survival (50). CPVL (Carboxypeptidase, Vitellogenic-Like) is an enzyme expressed in antigen-presenting cells that participates in antigen processing, thereby contributing to effective immune surveillance (51). GPR183 (G Protein-Coupled Receptor 183), also known as EBI2, is highly expressed on dendritic cells and regulates their positioning within tissues (52–54). Furthermore, GPR183 modulates immune responses by influencing interferon signaling, autophagy, and bacterial growth control during *Mycobacterium tuberculosis* infection (55).

### Characterization of Proliferating and Stromal Cell Populations in the Alveolar Microenvironment

Cluster 3 contains proliferating cells based on elevated expression of several key markers associated with active cell division, including MKI67, CIT, CENPF, and VIM (**Figure 5E**). These markers are characteristic of cells undergoing mitosis and cell cycle progression. MKI67 (Ki-67) is a nuclear protein expressed during all active phases of the cell cycle (G₁, S, G₂, and M phases) but absent in resting (G₀) cells (56), making it a well-established indicator of cellular proliferation (57). Notably, increased MKI67 expression has been reported in hypoxic lungs (58). CIT (Citron Rho-Interacting Serine/Threonine Kinase) plays a critical role in cytokinesis, the final stage of cell division, and is essential for proper neurodevelopment during embryogenesis (59,60). CENPF (Centromere Protein F) associates with kinetochores during mitosis and is vital for chromosome segregation and mitotic progression (61). VIM (Vimentin), an intermediate filament protein, is often upregulated in proliferating cells and during epithelial-mesenchymal transition, a process linked to increased cellular motility and proliferation (62,63).

In addition to these markers, Cluster 3 showed high expression of TOP2A (Topoisomerase II Alpha), an enzyme that modulates DNA topology during transcription and is crucial for DNA replication and cell division. TOP2A expression correlates with cellular proliferation and has been identified as a hub gene in various cancers, including lung tumors (64). IQGAP3 (IQ Motif Containing GTPase Activating Protein 3) facilitates cell cycle progression and is required for proper proliferation (65). Moreover, IQGAP3 is a downstream target of the YAP signaling pathway, which regulates organ size and tumorigenesis, further emphasizing the proliferative nature of this cluster (66).

Cluster 4 consists of epithelial cells and fibroblasts based on elevated expression of markers ALDH3A1, IGFBP5, CCDC187, TTC34, and CFAP100 (**Figure 5E**). These genes are commonly associated with epithelial identity and cellular stress responses. ALDH3A1 (Aldehyde Dehydrogenase 3 Family Member A1) is highly expressed in mammalian corneal epithelial cells and plays a critical role in protecting cells from UV radiation–induced damage and lipid peroxidation. Beyond its protective functions, ALDH3A1 also acts as a negative regulator of the cell cycle, influencing cellular proliferation (67). IGFBP5 (Insulin-Like Growth Factor Binding Protein 5) is upregulated in mammary epithelial cells, where it modulates insulin-like growth factor signaling, thereby impacting cell survival and apoptosis (68).

### Impact of Natural Aging on Lung Immune Cell Landscapes

To investigate the impact of natural aging on the lung immune cell landscape, we analyzed scRNA-seq data from AMs and other immune cell populations in young and old rhesus macaques. This analysis revealed distinct cellular distribution patterns and transcriptional changes associated with aging. Among the identified clusters, AMs were the most abundant cell type in both young and old groups.

To assess whether the proportions of immune cells differ significantly between young and old macaques, we performed z-tests for proportions. The analysis showed no statistically significant differences in the relative abundance of cell clusters during natural aging (**Figure 6A**). Furthermore, the spatial distribution of clusters on UMAP plots remained largely conserved between the two age groups (**Figure 6B**). To gain deeper insights into age-related pulmonary immune dynamics, we focused on transcriptional changes across all cell types, particularly alveolar macrophages. This approach allowed us to explore how natural aging influences specific immune cell functions while preserving overall cellular architecture and transcriptional landscape.

**Figure 6.**
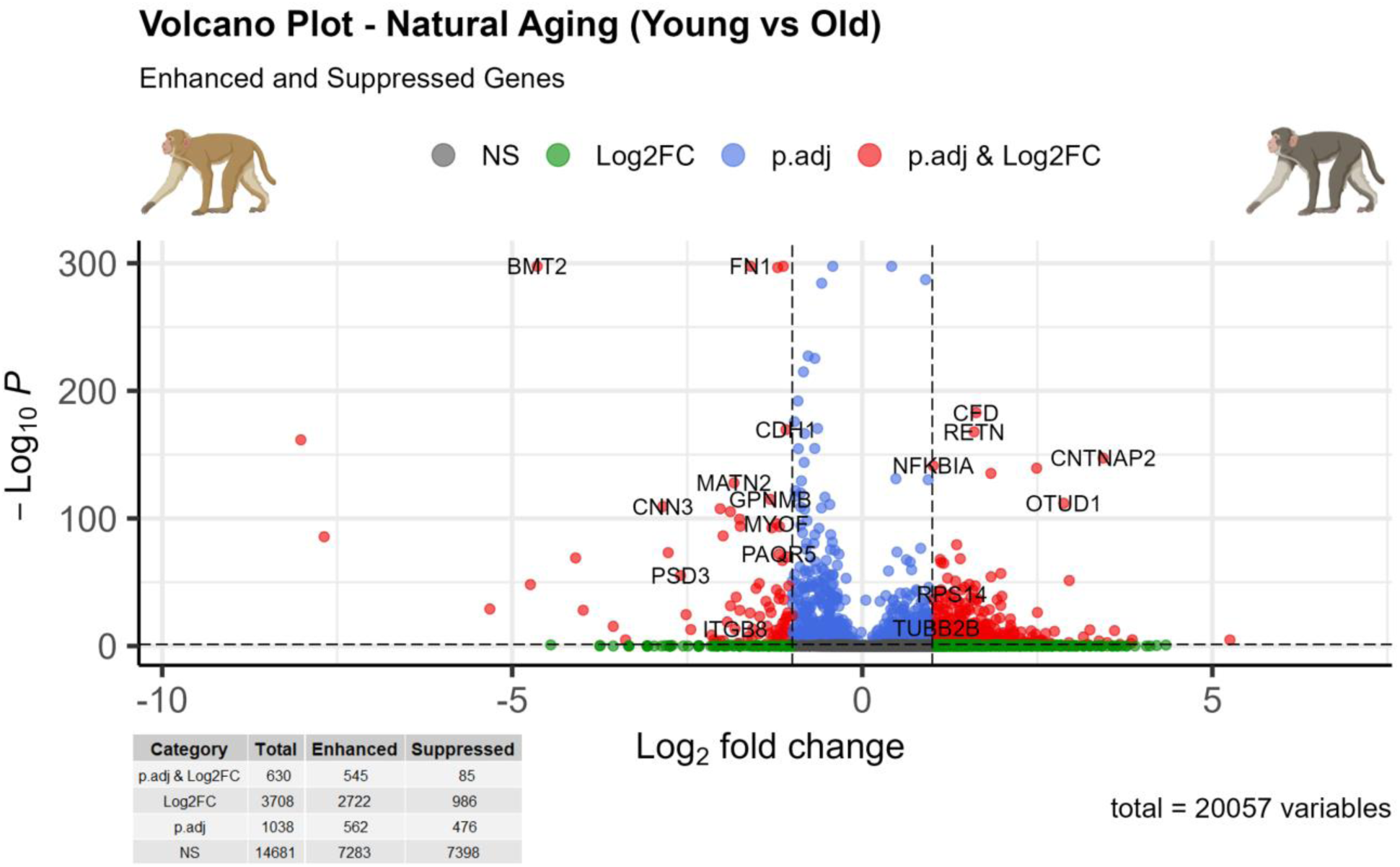
Upregulation of Gene Expression Characterizes Natural Aging in Alveolar Macrophage. Volcano plot showing differential gene expressions between young (n = 7) and old rhesus macaques (n = 5). Genes significantly downregulated in the old (< 0) and significantly upregulated in the old (> 0). Significantly regulated genes (red) are highlighted based on a threshold of adjusted *p-value* < 0.05 and log2 fold change (Log2 FC) > ±1.0. Upregulated genes dominate the transcriptional landscape, indicating overall gene enhancement in alveolar macrophages during natural aging.

### Enhanced Inflammatory and Metabolic Gene Expression in Aged Alveolar Macrophages

We examined differential gene expression profiles of AMs isolated from young and old rhesus macaques using scRNA-seq data. This analysis encompassed 20,057 genes, with significantly differentially expressed genes defined by a log₂ fold change (Log2 FC) greater than ±1.0 and an adjusted *p*-value less than 0.05 (**Figure 6**). Our results demonstrate robust transcriptional remodeling in AMs associated with natural aging. Consistent with prior studies, natural aging was linked to complex transcriptional changes (69), including the upregulation of genes involved in inflammation, immune responses, and lysosomal function (70). Notably, we observed an overall increase in gene expression in AMs from aged macaques compared to younger animals, underscoring the profound impact of aging on cellular transcriptional programs (**Figure 6**). This pattern may reflect compensatory responses to cellular stress or chronic immune activation.

Pathways associated with cytoskeletal organization, actin filament remodeling, cell adhesion, and wound healing were significantly downregulated in aged AMs (**Figure 7A–B**), suggesting a reduced capacity for tissue remodeling and repair. This decline may indicate impaired structural plasticity and compromised responsiveness to injury in the aging lung microenvironment. Key downregulated genes included BMX, LYZ, and CNN3, suggesting a decline in innate immune function and cytoskeletal remodeling with age (71–73).

**Figure 7.**
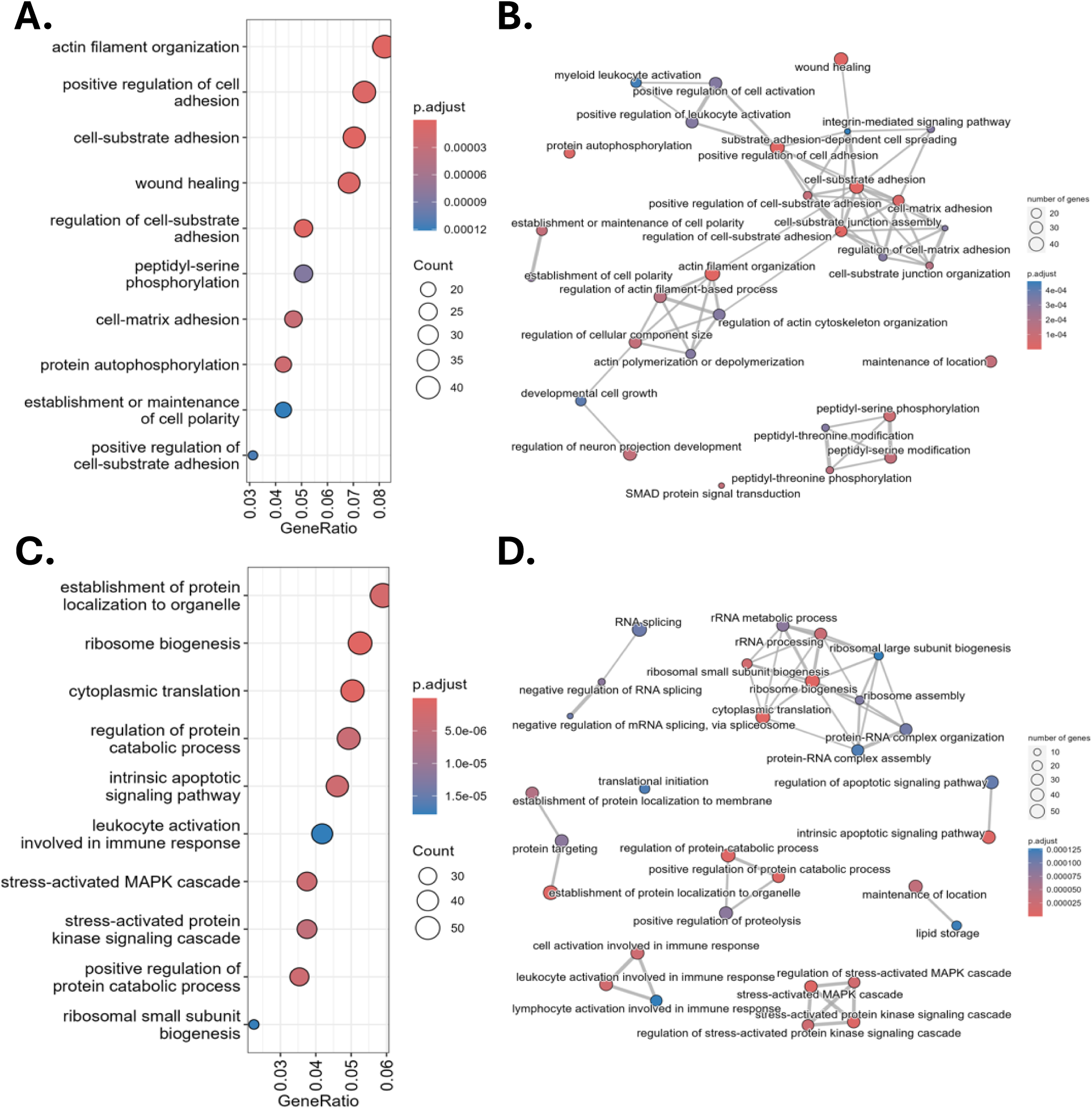
Age-Related Transcriptional Reprogramming in Alveolar Macrophages: Suppressed Protein Synthesis and Upregulated Structural Pathways. **(A & B) Pathways downregulated in old macaques**: Gene Ontology (GO) analysis reveals that aging is associated with decreased activation of structural and signaling pathways, including actin filament organization, cell adhesion, and wound healing, indicating suppressed cellular remodeling and compensatory mechanisms for tissue repair in response to aging-related stress. **(C & D) Pathways upregulated in old macaques**: enhanced pathways related to protein synthesis, including ribosome biogenesis, cytoplasmic translation, and RNA processing. These findings suggest an increase in cellular turnover and protein homeostasis in aging alveolar macrophages.

Conversely, genes encoding ribosomal proteins and regulators of protein synthesis, such as DDX3Y, RPS4Y1, RPLP2, and RPL13, were upregulated (**Figure 7C–D**), pointing to enhanced translational activity and increased demands for protein turnover, homeostasis and possible cellular stress responses (74). Additionally, lipid metabolism and immune regulatory genes including CFD, APOE, and APOC2 showed increased expression, consistent with metabolic reprogramming during aging (75,76).

Interestingly, transcripts with extreme fold changes, including uncharacterized genes ENSMMUG00000049951 and ENSMMUG00000028695, may represent novel markers or regulators of aging in alveolar macrophages. Functional enrichment analysis of these differentially expressed genes revealed pathways related to inflammatory responses, immune homeostasis, and translational regulation, aligning with documented age-related immune alterations (77–81).

### Suppressed Biological Processes in Aged Alveolar Macrophages Revealed by GO Analysis

To further understand how natural aging affects AM function, we performed Gene Ontology (GO) analysis using gene set enrichment analysis (GSEA) to identify significantly downregulated pathways in older rhesus macaques. This analysis revealed several suppressed biological processes, including ribosome biogenesis, cytoplasmic translation, and the establishment of protein localization to organelles (**Figure 7A**). These findings suggest a general decline in protein synthesis efficiency and intracellular trafficking, hallmarks of cellular aging.

In addition to core metabolic functions, the analysis also highlighted diminished activity in pathways associated with cellular turnover and stress response. Intrinsic apoptotic signaling and regulation of catabolic protein processes were both reduced in older animals, indicating impaired cellular homeostasis. Furthermore, pathways linked to immune function, including leukocyte activation and stress-activated MAPK cascades, were downregulated with age, pointing to a decline in immune responsiveness.

To contextualize these findings within broader biological networks, we applied a network-based GO analysis to visualize the interconnectedness of suppressed processes (**Figure 7B**). This approach reinforced the observation that essential regulatory pathways such as rRNA processing, RNA splicing, and translational initiation were significantly dampened in older macaques. The downregulation of processes like protein-RNA complex assembly, proteolysis regulation, and protein targeting further underscores a loss of transcriptional coordination and reduced capacity for adaptive responses in aged AMs.

### Transcriptional Rewiring Favors Tissue Repair Over Immunity in Aged Alveolar Macrophages

In AMs from aged rhesus macaques, GO enrichment analysis revealed a distinct upregulation of pathways involved in cytoskeletal organization, cell adhesion, and intracellular signaling (**Figure 7C**). Notably, pathways such as actin filament organization, positive regulation of cell adhesion, and cell-substrate adhesion were enriched, reflecting cytoskeletal remodeling and enhanced interaction with the extracellular matrix. These transcriptional changes suggest a shift toward cellular processes that support tissue integrity and repair in response to age-associated stressors. Enrichment of wound healing and cell-substrate adhesion pathways further underscores a potential compensatory response to ongoing or accumulated tissue damage in the aging lung. Additionally, elevated activity in phosphorylation-related pathways, including peptidyl-serine phosphorylation and protein autophosphorylation, points to modified intracellular signaling dynamics and regulatory adaptations during aging.

Network-based GO analysis reinforced these observations by highlighting highly interconnected biological processes enriched in aged macrophages (**Figure 7D**). Prominent among these were wound healing, positive regulation of leukocyte activation, and cell-matrix adhesion, indicating a combined response involving tissue repair and immune system modulation. Enhanced regulation of actin filament-based processes and junction assembly further emphasizes a broad reorganization of structural components, aligning with age-related remodeling of the pulmonary microenvironment (82). Phosphorylation-related pathways such as peptidyl-threonine phosphorylation suggest altered post-translational regulatory mechanisms as part of the aging process.

### Aging Alters Inflammatory and Metabolic Marker Expression in Alveolar Macrophages

To characterize the phenotypic shifts in AMs associated with natural aging, we examined the expression of genes previously linked to aging, immune regulation, and inflammation. Comparative analysis between young and old rhesus macaques revealed distinct transcriptional profiles in key regulatory genes (**Figure 8**). Several genes known to mediate immune signaling, antigen presentation, and metabolic regulation were upregulated in AMs from aged animals. These included *SIRT1*, *FOXO3*, *TP53*, *MTOR*, *KL*, *LYZ*, *FN1*, and *CHIT1*, each showing significant Log2 FC (highlighted in red). These genes are associated with critical cellular functions such as longevity signaling, phagocytosis, and extracellular matrix remodeling. The upregulation of these markers may reflect an adaptive mechanism by which aging AM attempt to preserve immune homeostasis in the pulmonary microenvironment. Additionally, *NFKBIA*, a gene encoding an inhibitor of the NF-κB pathway and a key modulator of inflammation and oxidative stress responses (83), was significantly upregulated in older macaques (shown in red). This increase indicates a possible tentative in control inflammation, but with a compromised ability to regulate inflammatory responses and maintain redox balance, potentially contributing to age-associated immune dysregulation.

**Figure 8.**
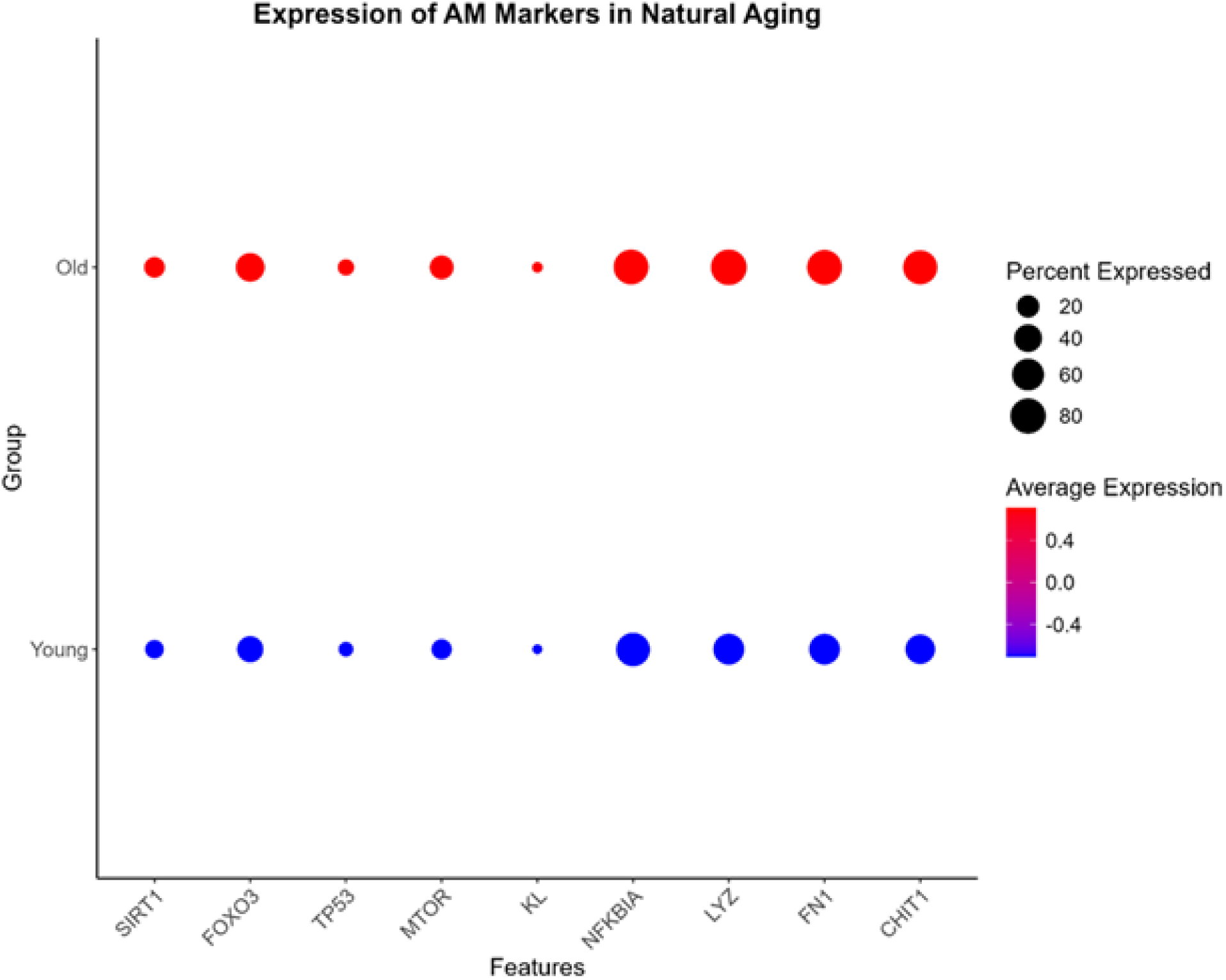
Expression of Aging and Inflammation Markers in Alveolar Macrophages During Natural Aging. Dot illustrates the expression levels of selected markers associated with aging and inflammation in alveolar macrophages from young and old rhesus macaques. The y-axis represents the group (young or old), and the x-axis lists the analyzed genes. The color gradient (blue to red) indicates the average expression levels, with red denoting higher expression and blue denoting lower expression. The size of the dots corresponds to the percentage of cells expressing each gene, with larger dots reflecting higher percentages.

## Discussion

This study demonstrates that aging in rhesus macaques is accompanied by heightened systemic inflammation and local transcriptional reprogramming of alveolar macrophages (AMs), a pattern that mirrors key aspects of immunosenescence observed in humans. By integrating plasma cytokine profiling with single-cell transcriptomics in alveolar macrophages of rhesus macaques, we demonstrate age-associated increased inflammatory markers on plasma and AMs in the lungs. Additionally, cytokine analysis of BAL supernatant demonstrated a different profile seen from plasma and AMs. Collectively, this data adds key information about the aging lung microenvironment.

Previous studies have reported age-associated shifts in immune cell populations, including significant declines in macrophage proportions across various tissues (84). Concurrently, increases in T cell populations have been observed in human lymphoid tissues during aging, suggesting compensatory immune remodeling (85).

Consistent with the concept of inflammaging, a chronic and low-grade systemic inflammation that increases with age (86–91), we observed significantly elevated plasma levels of pro-inflammatory cytokines, including IL-1β, TNF-α, and IL-6, in older macaques (**Figure 2**) (86,92,87,88,93,94,89–91). These cytokines, central mediators of innate immunity and tissue damage, contribute to age-related immune dysfunction (95,96). Their elevation supports the idea that aging induces a heightened basal immune tone, even in the absence of acute infection. The concurrent increase of IL-1RA, but unchanged IL-10 levels also suggests an incomplete or insufficient counter-regulatory response to inflammation (97).

The chemokine profile in aged plasma further supports sustained immune activation (**Figure 2**). Elevated plasma levels of MCP-1 (CCL2) and IL-8 (CXCL8) suggest increased recruitment of monocytes and neutrophils, respectively-hallmarks of a primed innate immune system. Increases in MIP-1α (CCL3) and MIP-1β (CCL4) also indicate broader leukocyte activation and trafficking, which may promote persistent inflammatory surveillance in peripheral tissues (98,99). Conversely, VEGFa, a critical factor for angiogenesis and tissue remodeling, was significantly reduced in aged plasma, aligning with previous findings of impaired tissue repair, fibrosis, and vascular dysfunction in aging (99–102).

Given the central role of macrophages in producing cytokines and orchestrating immune responses, we focused our investigation on alveolar macrophages, the dominant resident immune population in the lung alveolar space. AMs are both abundant and readily accessible via bronchoalveolar lavage (BAL), making them an ideal target for understanding age-associated immune changes in the lung microenvironment.

Using single-cell RNA sequencing, we found that aged AMs adopt a transcriptional profile that closely parallels the systemic inflammatory phenotype observed in plasma. Specifically, aged AMs upregulated IL1B, IL6, TNF, and CXCL8, mirroring the inflammatory mediators elevated in circulation (**Figure 3**). Notably, MIF (Macrophage Migration Inhibitory Factor) was consistently elevated across plasma, AMs, and BAL fluid samples (**Figure 9**). MIF is a pleiotropic cytokine with broad regulatory functions in inflammation, immune cell recruitment, and oxidative stress (103). Its upregulation across all compartments suggests that MIF may serve as a stable, systemic marker of immune aging in the lung. Given that MIF can be secreted rapidly without the need for de novo transcription and is involved in the induction of IL-6, TNF-α, IL-1β, and IL-8, its elevation may not only reflect, but also drive systemic and pulmonary inflammaging (92,93,104).

**Figure 9.**
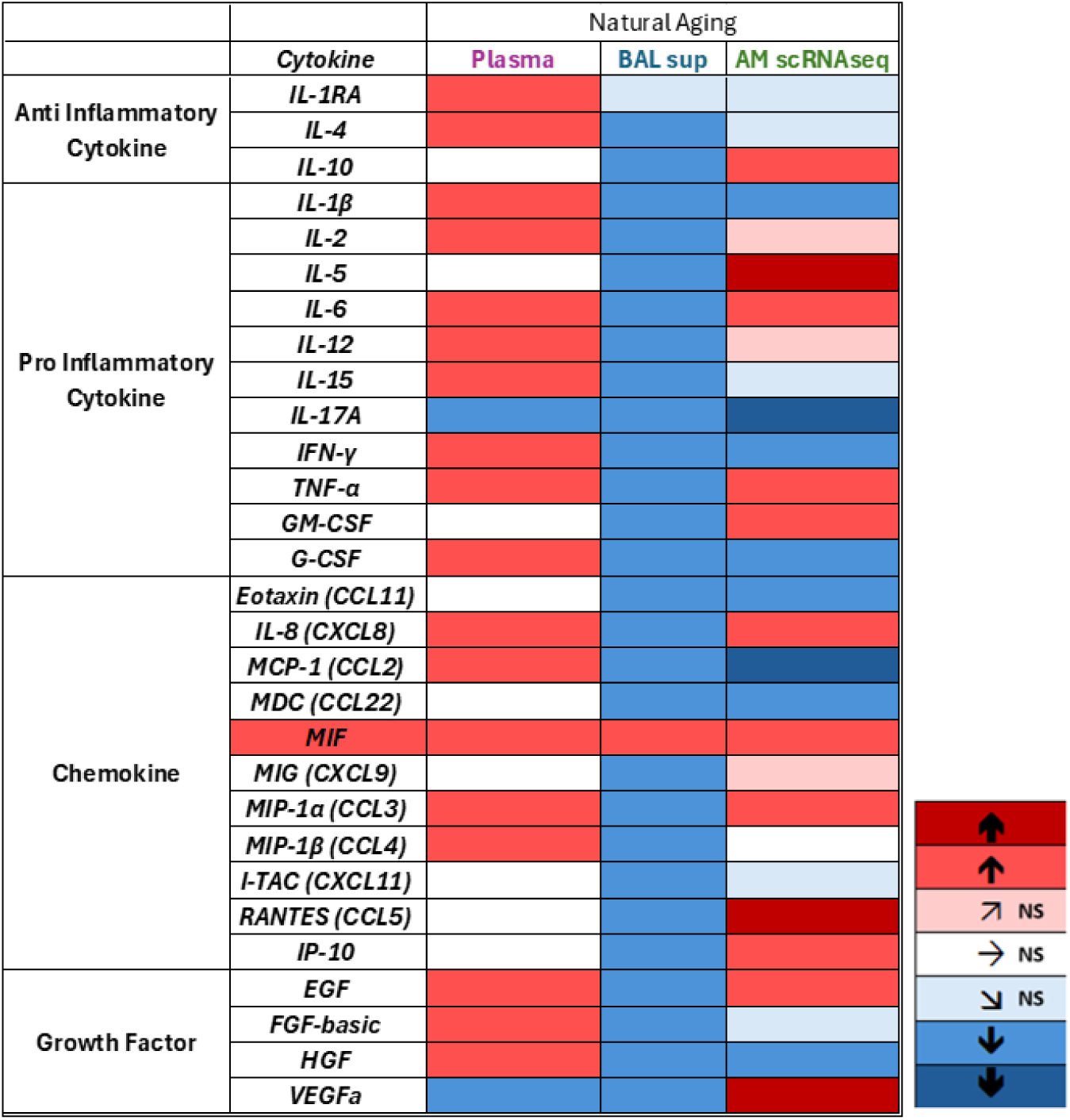
Heatmap of Cytokine Expression Highlights Pro-Inflammatory Shifts During Natural Aging. Heatmap summarizing the changes in cytokine expression across plasma, BAL supernatant, and alveolar macrophages (scRNA-seq) during natural aging. Pro-inflammatory cytokines (IL-6, TNF-α, IL-8 (CXCL8), and MIP-1α (CCL3), as well as growth factors (EGF, and HGF) are elevated in plasma and alveolar macrophages but show decreased levels in BAL supernatant. Notably, MIF is consistently elevated across all compartments, reinforcing its central role in age-associated inflammation. This compartment-specific regulation underscores the complexity of cytokine dynamics during aging.

The transcriptional landscape of aged AMs also revealed increased expression of genes associated with structural remodeling (*FN1*, *CHIT1*), immune regulation (*SIRT1*, *FOXO3*, *TP53*, *KL*), and phagocytosis (*LYZ*) (**Figure 8**). Upregulation of *DDHD1*, *TRPM2*, *APOE*, and *ACAD10* indicates metabolic reprogramming and lipid handling alterations, while markers such as *CD74*, *B2M*, and *SNORD89* point to enhanced antigen presentation and transcriptional regulation (**Figure 6**). These data suggest that aging AMs attempt to adapt to chronic inflammation by reinforcing metabolic, regulatory, and structural programs. In parallel, upregulation of NFKBIA, a key inhibitor of Nf-κB signaling, suggests tentative control over inflammation and redox balance, however concurrent with a weakened response, potentially contributing to uncontrolled immune activation (105).

GO and network-based enrichment analyses revealed a shift in transcriptional priorities in aged AMs (**Figure 7**). In aged AMs, pathways related to actin filament organization, cell adhesion, and wound healing were downregulated in aged AMs, indicating a reduction in cytoskeletal dynamics, cellular adhesion, and tissue repair activity. This suggests that aging is associated with suppressed remodeling capacity and a diminished ability to respond to tissue damage. In contrast, pathways related to ribosome biogenesis, cytoplasmic translation, and RNA processing were upregulated, highlighting an increase in protein synthesis and turnover. Together, these findings reveal extensive transcriptional reprogramming in AMs during natural aging. While younger animals exhibit dominant signatures of active protein synthesis and innate immune readiness, older individuals prioritize pathways associated with cell adhesion, cytoskeletal integrity, and wound healing. This shift likely reflects adaptive mechanisms aimed at maintaining homeostasis in the face of declining immune function and increased tissue stress. Such insights provide a nuanced understanding of how alveolar macrophages respond to the demands of aging in the lung.

Strikingly, BAL supernatant cytokine levels were not elevated with age, contrasting sharply with the pro-inflammatory plasma signature (**Figure 4**). This stands in contrast to both the plasma and AM transcriptional profiles, which showed a robust inflammatory signal. The reasons for this discrepancy remain unclear. One possibility is that while AMs represent the major immune cell population in the alveolar space, they may not be the sole contributors to the cytokine milieu measured in BAL fluid. Lung epithelial cells, which also produce cytokines and chemokines in response to environmental and immunologic cues, may undergo different age-related changes or exhibit diminished responsiveness. Additionally, BAL fluid represents a diluted compartment, and technical variables in lavage recovery could obscure low-level cytokine signals. Therefore, BAL cytokine measurements may not reliably reflect the true cellular activity occurring within the alveolar space.

In summary, our findings demonstrate that rhesus macaques exhibit a plasma cytokine profile in aging that closely resembles that of humans, supporting the translational relevance of this model. Alveolar macrophages in aged animals exhibit pro-inflammatory transcriptional reprogramming that mirrors systemic inflammaging, reinforcing their role as local amplifiers of immune aging. However, BAL supernatant cytokine measurements do not reflect these changes, pointing to either altered contributions from other lung-resident cells or limitations in the sampling method. These results underscore the importance of analyzing immune aging at the cellular and transcriptomic levels rather than relying solely on extracellular cytokine measurements in fluids.

### Implications and Future Directions

Collectively, our data highlight previously underappreciated roles for AMs as metabolically adaptive, but immunologically compromised cells in the aging lung. Our findings also highlight AMs as central mediators of pulmonary inflammaging. AMs in aged macaques exhibit transcriptional profiles consistent with chronic activation, metabolic adaptation, and impaired resolution of inflammation. This positions AMs as key amplifiers of pulmonary inflammaging and potential drivers of age-related lung vulnerability. Importantly, the divergence between the transcriptional inflammatory signature in AMs and the decreased cytokine levels in BAL supernatant highlights a critical disconnect between cellular activity and extracellular cytokine measurements. This discrepancy suggests the influence of other lung-resident cells, particularly epithelial cells, in shaping the local cytokine landscape, and emphasizes the need for compartment-specific analyses when studying immune aging. The consistent elevation of macrophage migration inhibitory factor (MIF) across plasma, AMs, and BAL fluid marks it as a promising biomarker and potential therapeutic target for age-associated inflammation. Given its capacity for rapid release and broad pro-inflammatory functions, targeting MIF signaling may offer a novel approach to attenuating chronic inflammation in the aging lung.

Future studies should investigate strategies to restore immune homeostasis and enhance tissue resilience in the aging lung. Key areas of interest include: the interplay between alveolar macrophages (AMs) and non-immune lung cells (such as epithelial and endothelial cells) in shaping the aging pulmonary microenvironment; the functional consequences of AM transcriptional reprogramming on pathogen clearance, tissue repair, and antigen presentation; and the therapeutic potential of targeting critical pathways identified in this study, such as MIF signaling, metabolic dysregulation, and impaired translational machinery, to mitigate inflammaging and promote healthy immune aging.

## Conclusion

This study establishes AMs as essential mediators of immune remodeling in the aging lung. Their long-lived nature, coupled with transcriptional reprogramming, positions them at the crossroads of inflammation, tissue repair, and immune senescence. By elucidating the molecular and compartmental complexity of lung aging, we provide a framework for understanding age-associated pulmonary decline and for developing targeted therapies aimed at restoring immune balance in elderly populations.

## Acknowledgements

We gratefully acknowledge the rhesus macaques that made this study possible. We thank the members of Kuroda Lab for their continual support and collaboration. We are especially grateful to: Lourdes Adamson, Andradi Villalobos, and Marcus Doyle for critical support in sample collection, experimental troubleshooting, and laboratory technique training; Tracy Rourke of the CNPRC Flow Cytometry Core for expert assistance with flow cytometry; Dr. Zhong-Min Ma for guidance during immunohistochemistry training; The Primate Assay Laboratory team, particularly Peter B. Nham, Amanda Carpenter, and Massiel Melendez, for technical expertise and daily support; The CNPRC Research Services and Pathology staff, especially Sarah Lockwood and Aaron Mark Allen, for their professionalism and efficiency during necropsy procedures; Also, thanks to Jennifer Watanabe for her kindness and steady encouragement during long hours at the bench.

## Funding

This research was supported by the National Institutes of Health (NIH) under the following grants:

- R01 MH107333 and R01 MH118139 (Woong-Ki Kim)
- R01 AI097059 and R33 AI110163 (Marcelo J. Kuroda)
- R01 HL139278 (Elizabeth S. Didier and Marcelo J. Kuroda)

Additional support was provided by:

- UC Davis School of Veterinary Medicine Office of Research & Graduate Education
- Comparative Medical Sciences Training Program (T32 OD011147)

## Conflicts of Interest

The authors declare no competing financial or non-financial interests.

## Author Contributions (with CRediT details)

**Conceptualization**: Jefferson G. C. Nagle; Marcelo J. Kuroda

**Methodology**: Jefferson G. C. Nagle, Raneesh Ramarapu, Laurent Zablocki

**Investigation**: Jefferson G. C. Nagle, Raneesh Ramarapu, Laurent Zablocki

**Data Curation**: Jefferson G. C. Nagle, Raneesh Ramarapu, Laurent Zablocki, Hai Duc Nguyen

**Formal Analysis**: Jefferson G. C. Nagle, Raneesh Ramarapu, Laurent Zablocki

**Visualization**: Jefferson G. C. Nagle, Raneesh Ramarapu, Laurent Zablocki

**Writing – Original Draft**: Jefferson G. C. Nagle; Hai Duc Ngueyn

**Writing – Review & Editing**: Marcelo J. Kuroda; Dennis Hartigan-O’Connor; Elizabeth S. Didier; Amir Ardeshir; Woong-Ki Kim

**Funding Acquisition**: Marcelo J. Kuroda; Elizabeth S. Didier; Woong-Ki Kim

**Supervision**: Marcelo J. Kuroda; Dennis Hartigan-O’Connor; Woong-Ki Kim

## Abbreviations

AMs: Alveolar macrophages
AAALAC: Association for Assessment and Accreditation of Laboratory Animal Care International
ACK: Ammonium-Chloride-Potassium
BAL: bronchoalveolar lavage
CNPRC: California National Primate Research Center
IACUC: Institutional Animal Care and Use Committee
PCA: Principal Component Analysis
NGS: Normal Goat Serum
UMAP: Uniform Manifold Approximation and Projection
scRNA-seq: single-cell RNA sequencing
SIV: Simian Immunodeficiency Virus
TNPRC: Tulane National Primate Research Center

## Notes

### Competing Interest Statement

The authors have declared no competing interest.

