## Supplemental Figures and Tables for "Characterization of immunosenescent alveolar macrophages in rhesus macaques"

**Supplemental Figure 1. Rhesus macaque lungs and lobes**

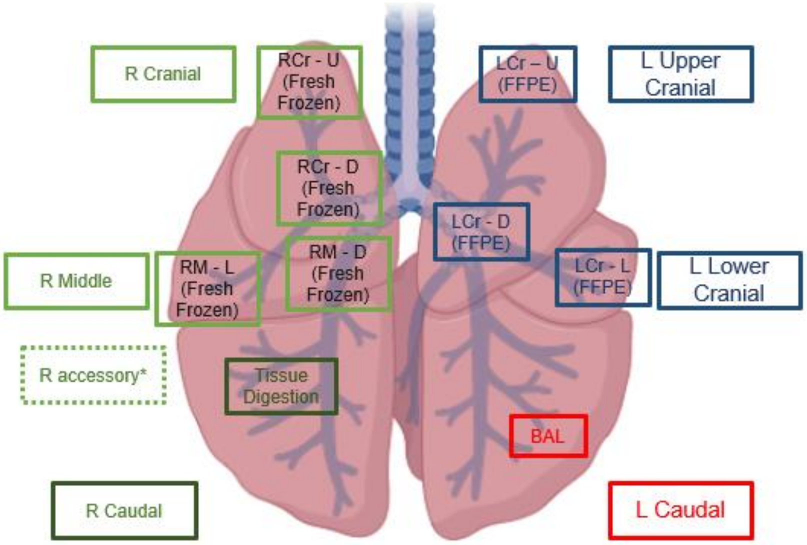


**Supplemental Figure 1.** Lung scheme with lobes and their usage in the project.

**Supplemental Figure 2. scRNA-seq Elbow Plot**

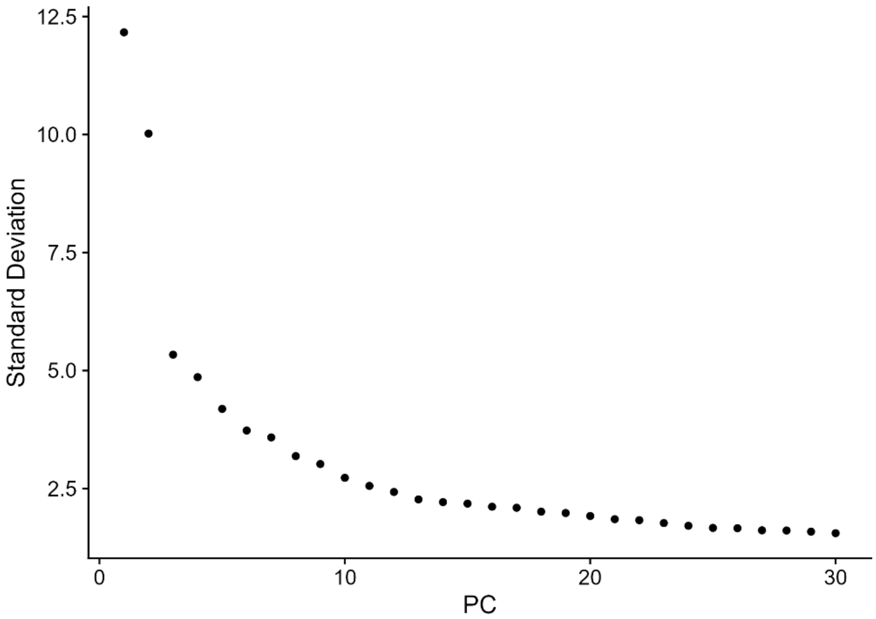


**Supplemental Figure 2.** The elbow plot shows a scatterplot with a curve that gradually decreases, illustrating the relationship between the principal components (PCs) and their corresponding standard deviations. The x-axis represents the PC numbers, while the y-axis indicates the standard deviations. The curve starts with a steep decline for the first few PCs, followed by gradual flattening, highlighting the transition point often referred to as the "elbow." This point marks the optimal number of PCs to retain for downstream analysis.

### Supplemental Figure 3. Clustree Analysis

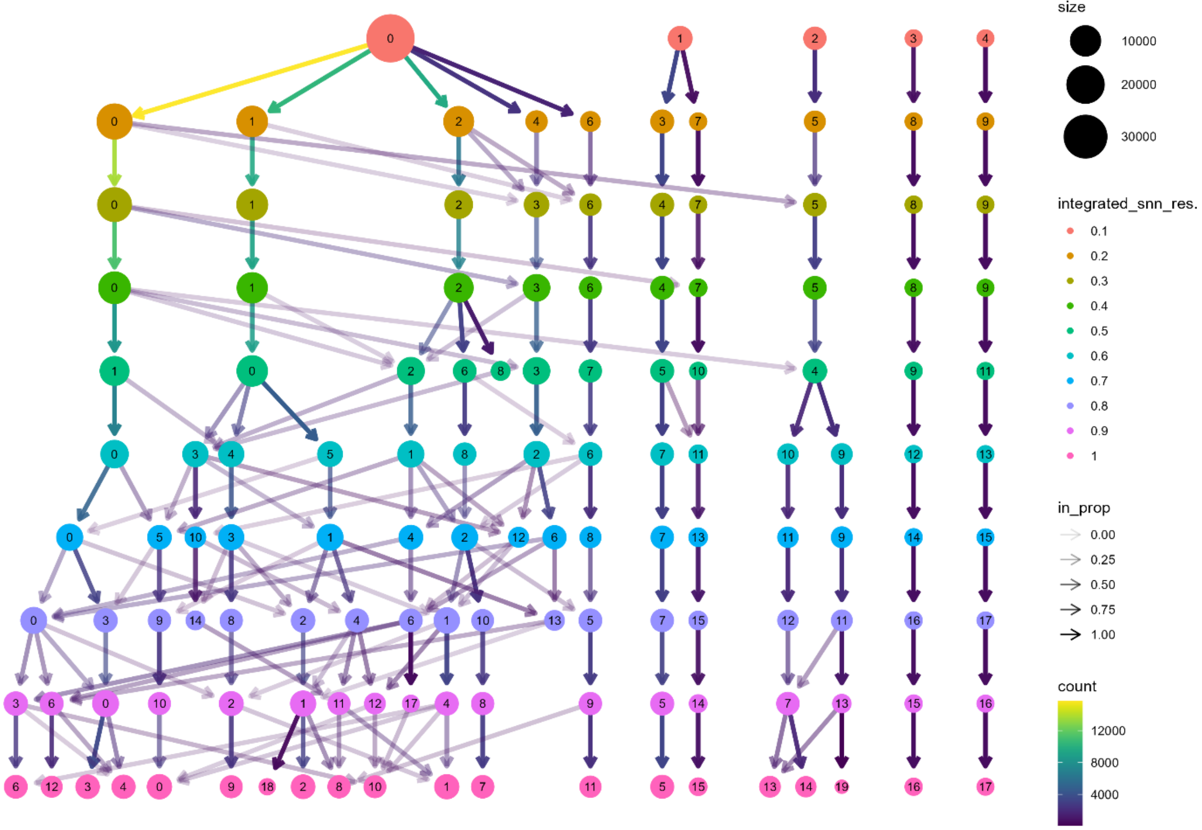


**Supplemental Figure 3.** *Clustree* analysis of cell isolates from BAL of 9 rhesus macaques using the 3000 most significant genes. At resolution of 0.2, we see the split of major clusters through the formation of branches, and transitions with increased resolutions. Because of that, resolution 0.1 offers a clear, interpretable clustering structure, balancing the trade-off between capturing cellular heterogeneity and maintaining meaningful biological groupings.

**Supplemental Table 1. Demographic Table of Natural Aging Plasma Study**

**Demographic Analysis of Natural Aging (Young Uninfected vs Old Uninfected)**

| Group | N | Mean (SD) | Median | Q1 - Q3 | Min - Max | Male (%) | Female (%) |
|---|---|---|---|---|---|---|---|
| Plasma | | | | | | | |
| Young | 36 | 8.9 (2.03) | 8.6 | 7.5 - 10.5 | 5.4 - 12.0 | 2.9 | 97.1 |
| Old | 69 | 18.9 (2.26) | 19.2 | 17.1 - 20.2 | 15.0 - 23.2 | 24.5 | 75.5 |
| BAL Sup | | | | | | | |
| Young | 26 | 6.5 (1.39) | 6.0 | 5.3 - 7.8 | 5.0 - 10.6 | 92.3 | 7.7 |
| Old | 13 | 19.1 (1.83) | 18.9 | 18.4 - 19.8 | 15.1 - 22.6 | 7.7 | 92.3 |

*Denominator based on number of non-missing responses.*

**Supplemental Table 2. Cytokine/chemokine analysis of plasma**

### Luminex Analysis of Cytokines in Plasma Samples

Young (5-12 years) vs Old (15-24 years)

| Cytokine | Young Median (Min - Max) | Old Median (Min - Max) | Exact p-value | p-value (adjusted) |
|---|---|---|---|---|
| IL-1RA | 30.785 (0.85 - 177) | 132.155 (28.85 - 935.2) | 0.000 | *** |
| IL-4 | 16.49 (2.75 - 64.42) | 29.01 (9.47 - 429) | 0.028 | * |
| IL-10 | 3.92 (0.99 - 10.07) | 3.645 (1 - 10.26) | 0.870 | NS |
| IL-1β | 6.19 (1.27 - 34.97) | 25.43 (15.42 - 48.46) | 0.000 | *** |
| IL-2 | 18.25 (4.28 - 234.11) | 173.445 (0.65 - 343.19) | 0.000 | *** |
| IL-5 | 1.2 (0.31 - 3.04) | 1.075 (0.12 - 2.55) | 0.258 | NS |
| IL-6 | 4.08 (0.39 - 15.93) | 6.59 (0.12 - 18.24) | 0.015 | * |
| IL-12 | 79.185 (12.63 - 1117.9) | 659.79 (160.7 - 1401.01) | 0.000 | *** |
| IL-15 | 7.75 (1.24 - 46.2) | 42.805 (18.99 - 75.71) | 0.000 | *** |
| IL-17A | 1.535 (0.07 - 4.35) | 0.26 (0.03 - 2.87) | 0.001 | ** |
| IFN-γ | 25.525 (2.8 - 67.77) | 34.36 (9.67 - 78.09) | 0.008 | * |
| TNF-α | 18.71 (0.07 - 83.41) | 51.47 (11.11 - 127.6) | 0.000 | *** |
| GM-CSF | 4.345 (0.2 - 12.63) | 4.63 (0.38 - 11.2) | 0.413 | NS |
| G-CSF | 52.43 (2.01 - 195.77) | 195.765 (47.67 - 279.05) | 0.000 | *** |
| Eotaxin (CCL11) | 411.56 (342.52 - 694.94) | 516.45 (152.67 - 1320.76) | 0.291 | NS |
| IL-8 (CXCL8) | 647.26 (13.68 - 6827.2) | 4065.355 (113.8 - 10000) | 0.001 | ** |
| MCP-1 (CCL2) | 234.7 (88.16 - 850.13) | 449.38 (151.12 - 1014.02) | 0.000 | *** |
| MDC (CCL22) | 727.11 (276.05 - 1538.5) | 552.1 (211.45 - 1581.8) | 0.271 | NS |
| MIF | 185.835 (61.2 - 535.09) | 659.28 (144.11 - 1304.47) | 0.002 | ** |
| MIG (CXCL9) | 115.32 (51.19 - 154.3) | 81.71 (40.33 - 151.39) | 0.078 | NS |
| MIP-1α (CCL3) | 25.93 (3.19 - 63.68) | 48.34 (5.09 - 85.39) | 0.000 | *** |
| MIP-1β (CCL4) | 19.05 (0.41 - 49.15) | 25.355 (4.41 - 50.03) | 0.005 | ** |
| I-TAC (CXCL11) | 23.11 (8.96 - 121.57) | 58.07 (5.39 - 222.41) | 0.293 | NS |
| RANTES (CCL5) | 10599.89 (4572.9 - 19187.9) | 7900.235 (2215.11 - 27746.3) | 0.392 | NS |
| EGF | 30.425 (15.9 - 43.98) | 56.635 (10.31 - 135.06) | 0.007 | * |
| FGF-basic | 15.69 (9.53 - 18.54) | 19.21 (8.83 - 35.43) | 0.033 | * |
| HGF | 120.86 (87.41 - 824.23) | 1340.215 (87.77 - 3752.31) | 0.022 | * |
| VEGFa | 113.19 (0.97 - 223.82) | 1.295 (0.02 - 5.48) | 0.000 | *** |

Adjusted p-values (FDR) significance shown.

**Supplemental Table 3. Cytokine/chemokine analysis of BAL supernatant.**

### Luminex Analysis of Cytokines in BAL Supernatant Samples

Young (5-12 years) vs Old (15-24 years)

| Cytokine | Young Median (Min - Max) | Old Median (Min - Max) | Exact p-value | p-value (adjusted) |
|---|---|---|---|---|
| IL-1RA | 181.78 (41.92 - 581.23) | 131.57 (79.68 - 520.86) | 0.894 | NS |
| IL-4 | 24.25 (8.95 - 52.84) | 7.075 (3.71 - 9.07) | 0.000 | *** |
| IL-10 | 3.6 (0.42 - 15.1) | 0.68 (0.26 - 1.13) | 0.000 | *** |
| IL-1β | 11.39 (1.68 - 34.28) | 2.66 (0.45 - 12.14) | 0.006 | ** |
| IL-2 | 169.505 (42.73 - 632.41) | 64.55 (15.67 - 96.4) | 0.000 | *** |
| IL-5 | 3.73 (1.27 - 13.42) | 1.69 (0.92 - 2.12) | 0.000 | *** |
| IL-6 | 4.21 (1.47 - 10.29) | 1.3 (0.46 - 2.9) | 0.000 | *** |
| IL-12 | 290.91 (78.4 - 935.2) | 106.99 (54.03 - 128.87) | 0.000 | *** |
| IL-15 | 135.18 (8.23 - 776.87) | 76.04 (33.86 - 170.03) | 0.020 | * |
| IL-17A | 25.485 (0.56 - 114.22) | 26.62 (0.56 - 31.32) | 0.035 | * |
| IFN-γ | 73.985 (3.72 - 251.89) | 7.35 (1.37 - 26.22) | 0.000 | *** |
| TNF-α | 50.38 (6.97 - 180.45) | 14.38 (1.04 - 23.72) | 0.000 | *** |
| GM-CSF | 8.02 (1.11 - 17.69) | 1.675 (0.27 - 3.26) | 0.000 | *** |
| G-CSF | 382.64 (52.11 - 1249.8) | 213.11 (41.08 - 462.53) | 0.014 | * |
| Eotaxin (CCL11) | 2.59 (0.23 - 9.18) | 0.45 (0.04 - 4.21) | 0.001 | ** |
| IL-8 (CXCL8) | 2.89 (2.36 - 4.17) | 0.8 (0.54 - 1.07) | 0.000 | *** |
| MCP-1 (CCL2) | 166.25 (48.1 - 782.59) | 60.67 (34.49 - 149.03) | 0.000 | *** |
| MDC (CCL22) | 1803.46 (465.76 - 4699.91) | 584.44 (318.78 - 719.21) | 0.000 | *** |
| MIF | 1869.28 (586.98 - 4039.06) | 3593.04 (866.62 - 13479.3) | 0.018 | * |
| MIG (CXCL9) | 33.86 (4.35 - 93.35) | 13.33 (5.7 - 34.77) | 0.000 | *** |
| MIP-1α (CCL3) | 51.93 (15.86 - 158.93) | 27.745 (6.78 - 33.7) | 0.000 | *** |
| MIP-1β (CCL4) | 72.68 (3.97 - 215.75) | 15.03 (3.81 - 43.09) | 0.000 | *** |
| I-TAC (CXCL11) | 47.03 (18.47 - 114.03) | 27.22 (14.95 - 48.77) | 0.000 | *** |
| RANTES (CCL5) | 18.52 (6.62 - 48.12) | 9.21 (2.79 - 27.56) | 0.007 | ** |
| IP-10 | 2.845 (1.07 - 6.02) | 1.25 (0.95 - 1.54) | 0.000 | *** |
| EGF | 7.3 (2.24 - 20.73) | 2.76 (0.48 - 7.27) | 0.000 | *** |
| FGF-basic | 11.46 (3.94 - 24.58) | 6.12 (2.92 - 7.69) | 0.000 | *** |
| HGF | 73.45 (14.88 - 181.33) | 20.395 (10.89 - 28.96) | 0.000 | *** |
| VEGFa | 12.835 (5.87 - 29.94) | 10.29 (3.97 - 13.98) | 0.004 | ** |

Adjusted p-values (FDR) significance shown.

**Supplemental Table 4. Silhouette Score Test for Dimensions and Resolutions**

| Dimensions | Resolution | Average Silhouette Score |
|---|---|---|
| 10 | 0.1 | 0.1631 |
| 10 | 0.2 | 0.1577 |
| 10 | 0.3 | 0.0718 |
| 15 | 0.1 | 0.1282 |
| 15 | 0.2 | 0.1178 |
| 15 | 0.3 | 0.0139 |
| 20 | 0.1 | 0.1092 |
| 20 | 0.2 | 0.0996 |
| 20 | 0.3 | 0.0107 |
| 30 | 0.1 | -0.0679 |
| 30 | 0.2 | -0.0341 |
| 30 | 0.3 | -0.0068 |

**Supplemental Table 5. Silhouette Score Test for Number of Genes**

| Threshold | Average Silhouette Score |
|---|---|
| 2000 | 0.1577 |
| 3000 | 0.1643 |
| 4000 | 0.1641 |
| 4500 | 0.1604 |
| 5000 | 0.1597 |
| 6000 | 0.1566 |
| 8000 | 0.1542 |
| 10000 | 0.1512 |
| 15000 | 0.1478 |

**Supplemental Table 6. Natural aging trend of cytokines analysis between plasma and BAL supernatant.**

| Cytokine Category | Cytokine | Plasma | BAL |
|---|---|---|---|
| Anti-inflammatory | *IL-1RA* | ↑ | NS |
| | *IL-4* | ↑ | ↓ |
| | *IL-10* | NS | ↓ |
| Pro-inflammatory | *IL-1β* | ↑ | ↓ |
| | *IL-2* | ↑ | ↓ |
| | *IL-5* | NS | ↓ |
| | *IL-6* | ↑ | ↓ |
| | *IL-12* | ↑ | ↓ |
| | *IL-15* | ↑ | ↓ |
| | *IL-17A* | ↓ | ↓ |
| | *IFN-γ* | ↑ | ↓ |
| | *TNF-α* | ↑ | ↓ |
| | *GM-CSF* | NS | ↓ |
| | *G-CSF* | ↑ | ↓ |
| Chemokine | *Eotaxin (CCL11)* | NS | ↓ |
| | *IL-8 (CXCL8)* | ↑ | ↓ |
| | *MCP-1 (CCL2)* | ↑ | ↓ |
| | *MDC (CCL22)* | NS | ↓ |
| | *MIF* | ↑ | ↑ |
| | *MIG (CXCL9)* | NS | ↓ |
| | *MIP-1α (CCL3)* | ↑ | ↓ |
| | *MIP-1β (CCL4)* | ↑ | ↓ |
| | *I-TAC (CXCL11)* | NS | ↓ |
| | *RANTES (CCL5)* | NS | ↓ |
| | *IP-10* | NA | ↓ |
| Growth Factor | *EGF* | ↑ | ↓ |
| | *FGF-basic* | ↑ | ↓ |
| | *HGF* | ↑ | ↓ |
| | *VEGFa* | ↓ | ↓ |
